# Programmable Surface Receptor-Mediated Cell Targeting with Designer Antibody-Routed Nanoparticles (DARTs)

**DOI:** 10.64898/2026.10.18.757718

**Authors:** Goldie V. Roth, Isabella R. Gengaro, Negar Ahmadian, Kewa Gao, David Wang, Shanxiu Xu, Michael Chavez, Paul Finn, Aijun Wang, Lei S. Qi

**Affiliations:** Department of Chemical Engineering, Stanford University, Stanford, CA 94305; Biotechnology Training Program, Stanford University, Stanford, CA 94305; Sarafan ChEM-H, Stanford University, Stanford, CA 94305; Center for Surgical Bioengineering, Department of Surgery, School of Medicine, University of California, Davis, Sacramento, CA 95817; Institute for Pediatric Regenerative Medicine, Shriners Hospitals for Children, Sacramento, CA 95817; Department of Biomedical Engineering, University of California, Davis, Davis, CA 95616; Department of Bioengineering, Stanford University, Stanford, CA 94305; Department of Biomedical Data Science, Stanford University, Stanford, CA 94305; Biohub, San Francisco, CA 94158

## Abstract

Precise *in vivo* delivery remains a massive barrier to therapeutic genome editing. Recently, major strides have been made towards cell type-specific delivery, with potential to improve the safety profiles of future gene therapies. However, programming such targeted tropism is limited by the minimal repertoire of scFv sequences, nanobodies, and viral fusogens that enable cell-type specificity, and the slow, iterative design-build-test cycle required to identify and optimize them. Here, we overcome this bottleneck by developing the <u>d</u>esigner <u>a</u>ntibody-routed (DART) platform: a modular, plug-and-play tool that decouples delivery vehicle production from tropism engineering. By presenting generic, bioorthogonal antibody-binding interfaces on the surfaces of enveloped particles, the DART system enables conjugation of those particles with off-the-shelf antibodies that direct cell entry without requiring genetic or chemical manipulation. <u>D</u>esigner <u>a</u>ntibody-routed virus-like particle<u>s</u> (DARTs) leverage this facile conjugation method to interrogate and identify transduction-competent surface antigens across cell types. They enable combinatorial targeting and the discovery of antigens that synergistically enhance transduction. Finally, DARTs function in the fetal murine nervous system, empirically identifying CD56 as a previously uncharacterized surface antigen enabling editing in hard-to-transduce neural populations. Overall, the DART platform provides a universal tool for rapidly profiling unique, targetable antigens and leveraging them for precision editing applications.

## Main

As genome editing tools become increasingly capable of treating diverse disease indications, the primary barrier to their widespread use remains the development of universal, programmable *in vivo* delivery methodologies^1–3^. Popular viral delivery tools have significant limitations: adeno-associated virus (AAV), while cell-type specific and generally safe, has a small packaging capacity and is difficult to rationally engineer^1,3–6^. Furthermore, while viral and non-viral vectors can broadly target specific organs, directing them away from certain tissues or exclusively to distinct cellular subtypes, particularly in complex tissues like the brain, remains a major challenge; for example, lipid nanoparticles (LNPs), despite the growing number of formulations capable of targeting specific organs, generally lack granular cell-type specificity and retain significant off-target delivery to the liver^1,5–9^. While these technologies are undoubtedly suited for certain genome editing applications, therapeutics for many diseases continue to remain out of reach due to our inability to bring editors safely into target cells with efficiency and specificity.

Recently, virus-like particles (VLPs) have demonstrated the ability to safely and transiently deliver gene, epigenome, base, and prime editors as ribonucleoprotein complexes (RNPs) or mRNAs^10–21^. However, similarly to other enveloped delivery vehicles, current strategies to direct their tropism typically rely on “hard-coding” single-chain variable fragments (scFvs), nanobodies, or viral fusogens into the surfaces of the particles^10–15,22–33^. While effective for well-characterized antigen targets, this method is both rigid and labor-intensive, requiring a new, iterative design-build-test cycle for every antigen being tested. It is also majorly limited by the unpredictability of protein folding on the nanoparticle’s surface, which may hinder presentation of functional surface-tethered moieties. Perhaps more importantly, this strategy is deeply constrained by the small catalog and diversity of existing scFvs, nanobodies, fusogens, or other binders, which produces an equally small space of targetable cell surface antigens. Since little is known *a priori* about which antigens best facilitate vector internalization and cell entry, these issues make it difficult to build and optimize novel delivery vehicles for targeting diverse cell types or tissues. Consequently, the majority of the cell-surface proteome remains unexplored for potential entry sites, and cell type-specific delivery is limited to a few applications with well-validated targets.

To overcome these limitations, we reasoned that developing fully programmable delivery vehicles would require decoupling the vehicle and its therapeutic payload from the targeting mechanism. One existing strategy for doing so directs enveloped delivery modalities with antibodies tethered to the particle’s surface via protein AG (pAG) or click chemistry^30^. While such work has expanded cell type-specific targeting capacity, the antibody attachment mechanism is either 1) not bio-orthogonal or 2) requires time-consuming synthetic chemistry to attach N-hydroxysuccinimide (NHS)-benzylguanine esters to antibodies, limiting *in vivo* application and throughput. As such, we identified the need for a rapid, fully bio-orthogonal, and widely-applicable technology to direct enveloped delivery vehicles in a cell type-specific manner.

In response, we developed the <u>d</u>esigner <u>a</u>ntibody-routed (DART) system: a modular platform that elevates enveloped nanoparticles from rigid delivery vehicles with hard-wired tropism to programmable search modalities for targeted cellular entry. Rather than genetically reprogramming a nanoparticle’s outer envelope for each new target, DART technology uses generic surface binders that readily conjugate to inexpensive, off-the-shelf commercial antibodies. We demonstrate that the DART platform can rapidly sample diverse antibody repertoires to identify optimal antigen targets across a wide range of cell types, including those resistant to standard lentiviral transduction. Furthermore, <u>d</u>esigner <u>a</u>ntibody-routed virus-like particle<u>s</u> (DARTs) enable combinatorial targeting, wherein multiple antibodies program synergistic targeting behavior, thereby cooperatively enhancing specificity and efficiency in mixed cell populations. Finally, because *in vitro* binding often fails to predict *in vivo* transduction capacity, we harness DARTs as an empirical *in vivo* antigen discovery platform. Given persistent challenges in targeting nanoparticles to neurons^34^, we applied DARTs directly to the murine fetal brain to discover optimal entry points for neural progenitor cells. Through this search, we identified CD56 as a functional cell surface antigen for targeted gene editing *in vivo*. By converting the complex process of nanoparticle retargeting into a simple, “plug-and-play” assembly process, the DART platform provides a generalizable framework for profiling, programming, and optimizing cell-type-specific delivery for therapeutic genome-editing applications.

## Results

### Engineering a universal interface for programmable, bio-orthogonal, antibody-dependent tropism

To overcome the rigidity of genetically hard-coding each vector’s tropism into its outer envelope, we sought to create a platform with a straightforward workflow that decoupled nanoparticle manufacturing from programming of targeting logic. We engineered the designer antibody-routed (DART) platform to be appended to a “blank slate” nanoparticle, enabling rapid conjugation with off-the-shelf antibodies to probe cell types of interest for optimal transduction-competent surface receptors. Our design was optimized around virus-like particles (VLPs) due to their inherent safety and potential for *in vivo* gene-editing applications. We foresaw broader application of the DART platform into other enveloped delivery vehicles, such as lentiviruses, as an additional goal.

To target specific cell-surface proteins of interest, our platform leverages inexpensive, off-the-shelf antibodies to confer tropism, as they are widely available and can bind to large antigen pools. Our design criteria mandated a modular architecture where users could: 1) produce a batch of off-the-shelf, naïve VLPs with no inherent tropism; 2) select commercial antibodies against candidate cell surface antigens with favorable differential expression in on- vs off-target cell types; and 3) program the VLPs’ targeting specificity via a brief incubation step prior to transduction (**Fig. 1a**). This process would rapidly identify which antigens best mediate VLP entry into a cell type of interest.

**Figure 1:**
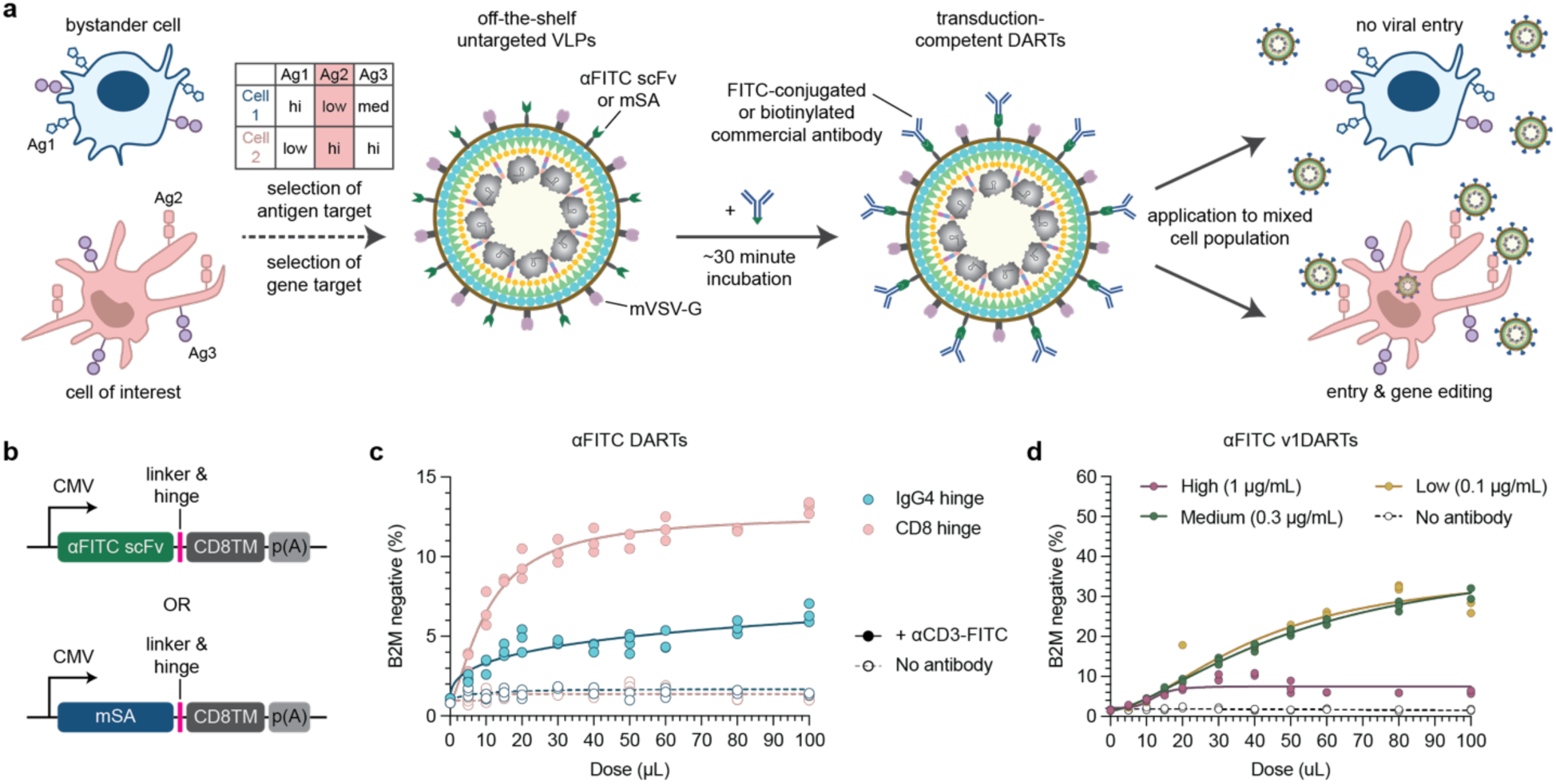
Generic surface-displayed ligands allow for antibody-dependent and specific genome editing. **a,** Schematic depicting the envisioned workflow for employing DARTs. Fusing an αFITC scFv or mSA to a transmembrane domain during VLP production creates particles decorated with either generic binder. Commercially available FITC-conjugated or biotinylated antibodies can then be attached to these viral particles via a brief incubation to confer the antibody’s targeting specificity. **b,** Depiction of the plasmid architecture of αFITC and mSA generic binders. **c,** For architecture optimization, DARTs containing sgB2M were produced with IgG4 and CD8α hinge variants connected to the αFITC scFv generic binder. Jurkat cells were either pre-treated with an αCD3-FITC targeting antibody or left unstained. The two groups were transduced with IgG4 and CD8α hinge architecture variants at a range of doses, and B2M knockout was quantified by flow cytometry 6 days later. DARTs were concentrated 5x prior to transduction. **d,** αFITC v1DARTs with the optimal CD8α hinge architecture (**c, pink**) were assessed at a range of doses. Prior to transduction, v1DARTs were incubated with four concentrations (0-1µg/mL) of αCD3-FITC targeting antibody for approximately 30 minutes. sgRNA sequence and concentration of DARTs was as in **c**. For **c**-**d**, all replicates are plotted and curves were fitted via sigmoidal four-parameter logistic regression. n=3 technical replicates per condition for all experiments. In **d**, two total outliers were identified via ROUT test (false discovery rate of 5%) and subsequently removed from analysis. Antibody clones used for transduction: αCD3, HIT3a. Antibody clone used for staining: αB2M, A17082A. Abbreviations: αFITC scFv, anti-FITC single-chain variable fragment; mSA, monomeric streptavidin; CD8TM, CD8α transmembrane domain; p(A), poly-A tail; B2M, β-2 microglobulin.

We engineered the DART platform by co-displaying two distinct functional moieties on the VLP envelope: 1) a fusion-competent but binding-deficient mutant VSV-G (mVSV-G) to facilitate endosomal escape^26,35^, and 2) a generic “capture” binder to anchor targeting antibodies. The former facilitates endosomal escape after particle internalization, while the latter confers tropism only when conjugated with a specific antibody targeter. Since lentiviruses and VLPs bud off the plasma membranes of producer cells, the external faces of these particles are decorated with our generic binders by expressing them fused to a transmembrane domain^10,12,13,15,24–27,29,30^.

For the capture binder, we first identified the ubiquity of fluorescein isothiocyanate (FITC) and biotin conjugations to commercial antibodies. We therefore chose an anti-FITC single-chain variable fragment (αFITC scFv; K_d_ = 0.75 nM^36,37^) and monomeric streptavidin (mSA; K_d_ = 2.8 nM^38^) as our universal ligands. To optimize surface display and membrane stability, we anchored these moieties to our nanoparticle envelopes via a CD8α transmembrane domain^10,12,13,24–26,39,40^ and evaluated both a CD8α hinge and a stabilized S228P IgG4 hinge to determine optimal capture binder architecture (**Fig. 1b**)^41,42^.

We tested these designs using Cas9-packaging, HIV-1-derived VLPs programmed to knock out the β-2 microglobulin (B2M) gene in Jurkat T lymphocytes. We transfected each of our four capture binder constructs into HEK293T producer cells alongside three other packaging plasmids; these expressed mVSV-G and engineered VLP components for packaging Cas9 RNP complexed with a single guide RNA (sgRNA) for B2M knockout. The resulting αFITC- or mSA-decorated <u>d</u>esigner <u>a</u>ntibody-routed virus-like particle<u>s</u> (DARTs) were applied at varying doses to cells pre-labeled with an αCD3 targeting antibody for 30 minutes, with excess antibody removed prior to transduction. CD3, a pan-T cell marker associated with the T cell receptor (TCR) complex, was selected as our first antigen target due to its high expression on Jurkat cells and precedent use in T cell-specific targeting^10,12,43^.

The CD8α hinge showed superior activity to the IgG4 hinge in both the αFITC and mSA DARTs (**Fig. 1c**, **SI Fig. 1a**), indicating the importance of this linker region. For both VLP formulations, B2M knockout with the CD8α hinge architecture was approximately 40-50% greater at the maximum dose. Notably, B2M editing was also negligible in the absence of FITC-conjugated or biotinylated αCD3 antibody. These findings designated the CD8α hinge as the superior architecture and demonstrated that DARTs were naïve without the addition of a targeting antibody, as desired. As such, we proceeded to label this CD8α hinge configuration as αFITC and mSA version 1 DARTs (v1DARTs).

It is worth noting the lower overall potency of mSA v1DARTs compared with the *α*FITC formulation. This finding may be explained by transduction interference from endogenous biotin or by the higher K_d_ of mSA. As a control for all future experiments, untransduced Jurkat cells were pre-incubated with the targeting antibody on the day of transduction to rule out any antibody-dependent B2M changes independent of DARTs’ activity; no effect was observed (**SI Fig. 1b**).

To identify the optimal hinge for v1DARTs, we initially stained target cells with FITC-conjugated or biotinylated antibodies, washed away the unbound fraction, and then added DARTs; we refer to this as the cell “pre-staining” method. However, for a more practical and relevant approach to antigen screening, we subsequently incubated candidate antibodies with DARTs for 30 minutes, then added the resulting mixture to target cells or administered it *in vivo*. This approach allowed targeting antibodies to be tested without first perturbing the cells.

With this new protocol, we hypothesized that the DART system would be sensitive to the concentration of targeting antibody used during this incubation. Too low of a concentration would insufficiently coat the generic capture binders, while a large excess could reduce overall transduction levels via a potential “Hook effect,” wherein free, non-conjugated antibody would bind target antigens and block transduction-competent DARTs from being endocytosed^44^. To understand this regime and thereby facilitate a more high-throughput workflow for our antigen screening applications, we characterized the antibody concentration dependence of αFITC v1DARTs within our Jurkat cell context.

We incubated αFITC v1DARTs – carrying Cas9 and an sgRNA for B2M knockout, as in **Fig. 1c** – with three concentrations of αCD3-FITC antibody for 30 minutes, then applied them to Jurkat cells. Low (0.1μg/mL) and medium (0.3μg/mL) antibody concentrations created similar dose-response curves, while clear inhibition was observed in the high (1μg/mL) condition (**Fig. 1d**). These findings outlined general guidelines for *in vitro* DARTs-antibody assembly and confirmed our suspicions regarding the use of excessive antibody concentrations. Importantly, regardless of nanoparticle dose, v1DARTs did not generate B2M knockout in the absence of targeter.

Notably, outside of inhibitory antibody concentrations, v1DARTs performed better when incubated directly with the targeting antibody (**Fig. 1d**), rather than added in a naïve state to pre-stained Jurkat cells (**Fig. 1c**). One rationale for this finding may be that the pre-stained cells prematurely internalize the antibody prior to DARTs’ attachment, thereby reducing transduction efficiency. Another explanation may be that the overall tripartite binding kinetics are less favorable with cell-antibody conjugates than DARTs-antibody conjugates.

To benchmark our non-covalent assembly strategy against traditional membrane-integrated engineering, we compared both αFITC and mSA v1DARTs to VLPs directed by an αCD3 scFv (αCD3 VLPs) with parallel architecture (**SI Fig. 1c**). Using our pre-staining method (as in **Fig. 1c**), we found that v1DARTs exhibited lower absolute editing efficiency compared to the αCD3 VLP control (**SI Fig. 1d**). This reduced efficacy may be due in part to our use of the lower-efficiency pre-staining strategy. Notably, this trend persisted even when varying the amount of generic capture binder on the surface of the DARTs (**SI Fig. 1e**), suggesting that the external interface was unlikely to be the cause of the discrepancy in editing rates. Quantification of Cas9 content within αFITC v1DARTs vs αCD3 VLPs via ELISA revealed that the latter packaged approximately 7.7 times more RNPs per particle (**SI Fig. 1f**), likely accounting for the differences in efficiency. The Cas9 encapsulation and overall editing rate gap between DARTs and αCD3 VLPs inspired further attention towards optimization of DARTs’ internal packaging and processing of the Cas9 RNP payload.

### Rational engineering of cargo-to-structure stoichiometry to enhance DART potency

Having established the basic parameters of our external targeting strategy, we next sought to maximize DARTs’ per-particle potency. VLP assembly is thought to depend on the stoichiometric balance between the endogenous, structural Gag-Pol polyprotein^45^, which is crucial for the formation and processing of the particle’s internal membrane, and the cargo-bearing Gag fusion protein, which packages the intended payload into the particle^10,11,16–18^. DARTs are produced by transfecting producer cells with four plasmids: plasmids A and B regulate the external surface of the particles, encoding mVSV-G and the generic capture binder, respectively; plasmid C expresses an sgRNA and the endogenous HIV-1 Gag-Pol polyprotein; and plasmid D expresses an sgRNA and a C-terminal truncated Gag fused to cargo (Cas9) via a cleavable linker (**Fig. 2a,b**). In this configuration, cargoes are physically bound to the VLP’s inner membrane, which forms around the payload prior to budding. As the VLP matures, the endogenous HIV-1 protease, expressed from plasmid C, separates the cargo from Gag by cutting at the protease recognition sequence contained within the cleavable linker (**SI Fig. 1g**).

**Figure 2:**
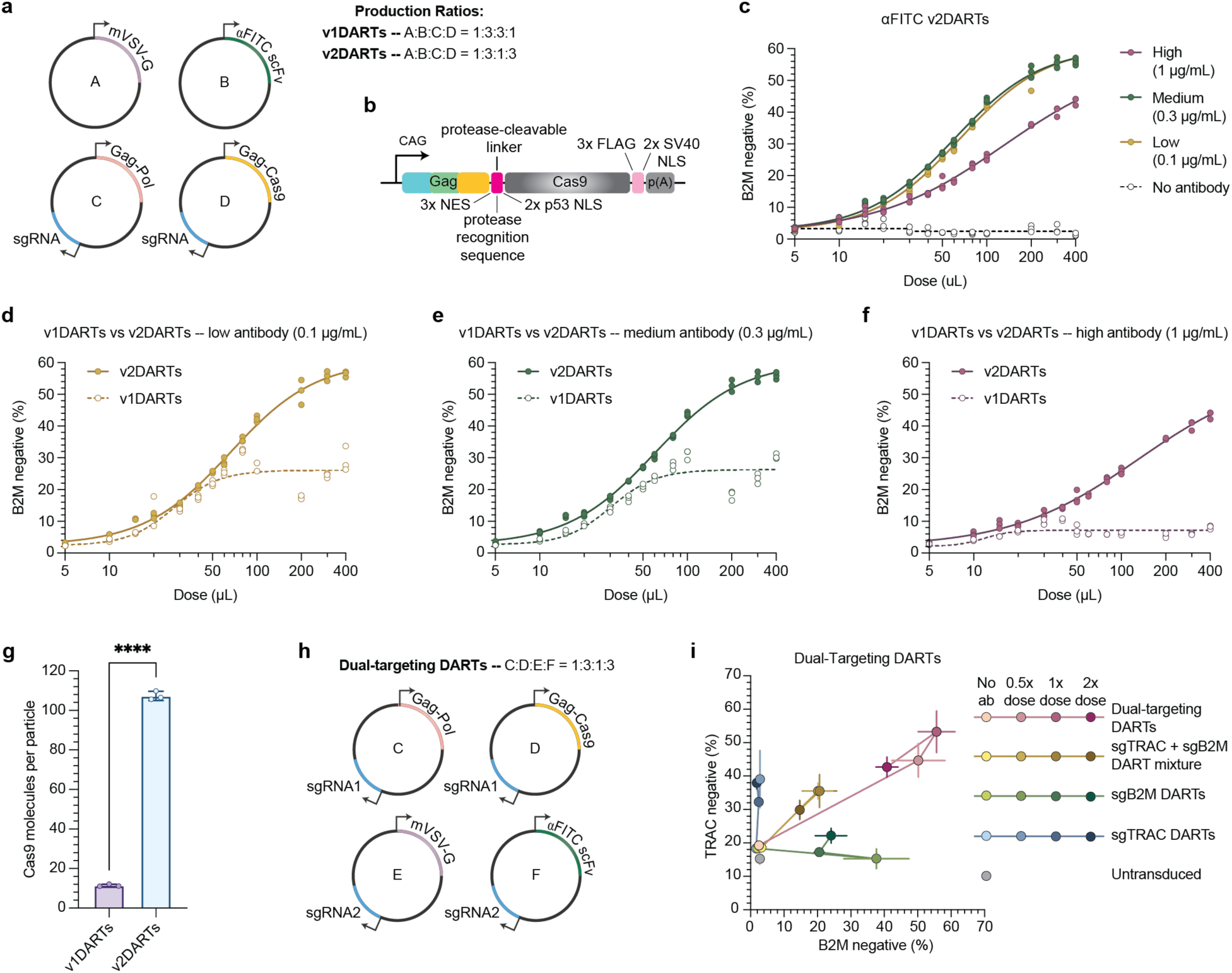
Modulation of DARTs’ internal composition improves gene editing efficiency and allows for simultaneous KO of multiple genes. **a,** Schematic of 4 plasmids used to produce DARTs and their transfection ratios for v1 vs v2 systems. **b,** Detailed diagram of the architecture of the Gag-Cas9 expression cassette. **c,** αFITC v2DARTs containing sgB2M were incubated with four concentrations (0-1µg/mL) of αCD3-FITC targeting antibody, then added to Jurkat cells in varying amounts to produce a dose curve. DARTs were concentrated 5x prior to transduction, and B2M knockout was quantified via flow cytometry 6 days post-transduction. **d-f**, Isolated comparisons of αFITC v1 and v2DARTs dose curves at low (**d**), medium (**e**), and high (**f**) targeting antibody concentrations. v2DARTs data is repeated from **c**. For **c-f**, all replicates are plotted and curves were fitted via sigmoidal four-parameter logistic regression. **g,** Quantification of Cas9 molecules per particle in αFITC v1 and v2DARTs. Cas9 was quantified by ELISA and normalized to particle concentration obtained via nanoparticle tracking analysis (NTA). Statistical comparison was performed via Welch’s t test; P<0.0001. **h,** Schematic of 4 plasmids used to produce dual-targeting DARTs and their transfection ratios for each system. Plasmids A and B from **a** were replaced with E and F. **i,** Flow cytometry-based quantification of simultaneous TRAC and B2M knockout imparted by DARTs carrying sgTRAC alone (blue), sgB2M alone (green), a 1:1 mixture of those two formulations (yellow), or dual-targeting DARTs packaging both sgRNAs simultaneously (pink). The untransduced condition is shown in gray. Increasingly dark shade in each color indicates increasing dose of DARTs, which were concentrated 40x prior to transduction. Jurkat cells were pre-stained with αCD7-FITC, and readout was quantified 6 days later. For all data, n=3 technical replicates per condition. Where error bars are plotted, data are all mean ± s.d. In **c**, one outlier was identified via ROUT test (false discovery rate of 5%) and subsequently removed from analysis. Antibody clones used for transduction: αCD3, HIT3a, αCD7, CD7-6B7. Antibody clones used for staining: αB2M, A17082A; TRAC, IP26. Abbreviations: NES, nuclear export signal; NLS, nuclear localization signal; B2M, β-2 microglobulin.

As mentioned above, a critical challenge in VLP design is optimizing the ratio between full-length Gag-Pol versus truncated Gag fused to cargo. It has been demonstrated that decreasing the ratio of these components can improve VLP function, though efficacy does not always scale directly with the amount of packaged RNP^11,17^. While more Gag-cargo equates to more effector molecules, an excessive amount may sterically hinder particle formation; furthermore, an insufficient concentration of full-length Gag-Pol can compromise particle maturation and structural integrity. As a result, standard VLP formulations often start by prioritizing structural stability, beginning optimization with a conservative ratio of Gag-Pol to Gag-cargo^10,11,17,18^.

Accordingly, we implemented a ratio of 3:1 Gag-Pol to Gag-Cas9 for v1DARTs. However, given v1DARTs’ relatively low efficacy and Cas9 encapsulation rate (**SI Fig. 1f**), we hypothesized that a more optimized VLP formulation could leverage increased cargo packaging. To test this hypothesis, we produced version 2 DARTs (v2DARTs) with an inverted Gag-Pol to Gag-Cas9 ratio of 1:3, which we reasoned would package roughly three times more editor molecules (**Fig. 2a**).

To assess the new cargo-to-structure stoichiometry, αFITC v2DARTs carrying the B2M knockout sgRNA were incubated with three concentrations of αCD3-FITC antibody, then applied to Jurkat cells to produce a dose curve (**Fig. 2c**). Across antibody concentrations, v2DARTs possessed higher potency at all doses and achieved a higher level of knockout at saturation than v1DARTs (**Fig. 2d-f**). Quantification of Cas9 contents via ELISA indicated that v2DARTs packaged over nine times more Cas9 protein than v1DARTs, with an average of 107 Cas9 molecules per particle (**Fig. 2g**). This ratio exceeded our initial hypothesis and explained the significant improvement in editing rates across the two versions. Notably, both v1 and v2DARTs produced negligible editing in the absence of targeting antibodies (**Fig. 1d, 2c**). Collectively, these findings suggest that the limiting factor for v1DARTs’ activity was the number of Cas9 molecules per VLP. By rationally tuning the internal packaging parameters, we produced a high-density particle that improves the probability of successful genome editing outcomes per entry event.

### DART engineering enables sgRNA co-encapsulation for efficient, multiplexed genome editing

Cell engineering often necessitates the simultaneous disruption of multiple genetic loci. Achieving this feat using a mixture of viruses is typically inefficient, due to the stochastic nature of co-transduction. To overcome this issue, we engineered DARTs to perform all-in-one, multiplexed editing at two genomic loci.

We first re-configured the modular, four-plasmid assembly to boost sgRNA delivery. Rather than introducing additional plasmids, which would likely diminish transfection and co-expression efficiency, we embedded secondary sgRNA expression cassettes directly into the plasmids encoding VLP surface proteins (mVSV-G and the generic capture binder) (**Fig. 2h**). Our design ensures that producer cells express and package both sgRNAs in approximately equivalent amounts, eliminating the need for viral mixtures.

We validated the resulting dual-targeting DARTs by programming them to simultaneously knock out T cell receptor alpha constant (TRAC) and B2M in Jurkat cells. We compared them against v2DARTs containing sgTRAC alone, sgB2M alone, or a mixture of the two VLP formulations. Dual-targeting DARTs showed significantly higher rates of double gene knockout than the combination of single-targeting DARTs when tested on Jurkat cells pre-stained with an αCD7-FITC antibody (**Fig. 2i**). They also outperformed both single-targeting formulations in knocking out their respective genes, suggesting that dual-targeting DARTs may package altogether more functional Cas9 RNP. Overall, these results indicate that DARTs can be simply and rationally engineered for multi-gene targeting.

#### Rational tuning of protease kinetics and nuclear import of cargo drives maximum potency

HIV-1-based VLPs frequently exhibit incomplete maturation and cleavage of the Gag-cargo linker due to limited proteolytic processing^10,14,16^. We confirmed this finding by analyzing v2DARTs via cryogenic transmission electron microscopy (cryo-EM), in which we observed only one particle with a processed, mature inner membrane; instead, 68.6% contained an immature inner membrane, and 30.7% were undefined (**Fig. 3a-c, SI Fig. 2a-c**). We surmised that this incomplete proteolytic processing may be causing a cargo-release issue, in which active Cas9 remained physically tethered to the viral structural lattice, hindering its diffusion to the nucleus following cellular entry and endosomal escape.

**Figure 3:**
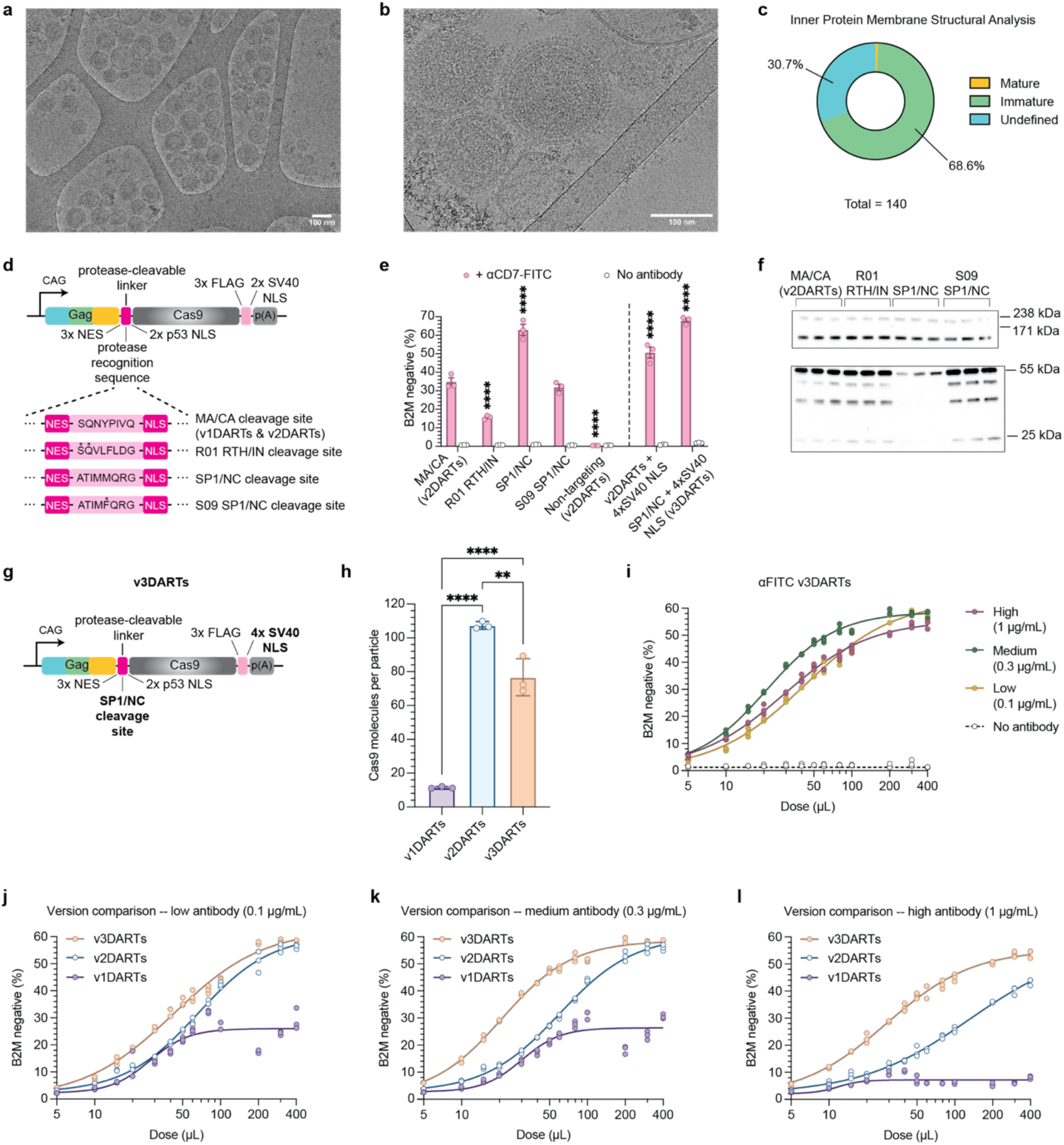
Rational optimization of DARTs’ structural components further improves gene editing outcomes. **a,b,** Cryo-EM images of αFITC v2DARTs. Scale bars = 100nm. **c,** Quantification of the inner protein membrane maturation status of 140 v2DARTs. For examples of particles with mature, immature, and undefined inner membranes, see **SI Fig. 2a-c**. **d,** The protease recognition sequence from v1 and v2DARTs (MA/CA) was replaced with three alternatives (below). Asterisks indicate residues mutated from the native HIV-1 sequence. **e,** B2M knockout in Jurkat cells induced by αFITC DARTs containing the four protease recognition sequences shown in **d**. Each formulation is named for the recognition sequence it contains. Formulations to the left of the dotted line contain two SV40 NLSs (as is standard in v1 and v2DARTs), while formulations to the right contain four. Non-targeting sgRNA targeted the genetic sequence for green fluorescent protein (sgGFP). A 1x dose of DARTs, defined as the amount of VLP generated in 2mL of producer cell media, was used for all conditions. Illustrated statistical significance represents P < 0.0001 via two-way ANOVA, with all groups compared to MA/CA (v2DARTs). **f,** Western blot comparing v2DARTs (lanes 1-3) to DARTs containing the three protease-cleavable linker variants depicted in **d** (R01 RTH/IN, lanes 4-6; SP1/NC, lanes 7-9; S09 SP1/NC, lanes 10-12). Molecular weights are shown to the right. **g,** Detailed diagram of the final Gag-Cas9 construct architecture for v3DARTs; changes from v2DARTs in **Fig. 2b** are highlighted in bold. **h**, Quantification of Cas9 molecules per particle in αFITC v1, v2, and v3DARTs, as in **Fig. 2g**. *, P<0.05; **, P<0.01; ***, P<0.001; ****, P<0.0001. **i**, A dose curve for αFITC v3DARTs was produced as in **Fig. 2c**. **j-l**, Isolated comparisons of αFITC v1, v2, and v3DARTs at low (**j**), medium (**k**), and high (**l**) targeting antibody concentrations. v3DARTs data is repeated from **i**. For **i-l**, all replicates are plotted and curves were fitted via sigmoidal four-parameter logistic regression. For all data, n=3 technical replicates per condition. Antibody clones used for transduction: αCD3, HIT3a, αCD7, CD7-6B7. Antibody clones used for staining: αB2M, A17082A. Abbreviations: MA/CA, matrix/capsid; RTH/IN, RNase H/Integrase; SP1/NC, spacer peptide 1/nucleocapsid; NES, nuclear export signal; NLS, nuclear localization signal; B2M, β-2 microglobulin.

To approach our hypothesis and contend with this proposed intrinsic barrier, we applied rational protein engineering to the protease-cleavable linker connecting Gag and Cas9 (**Fig. 2b**). Endogenous HIV-1 has ten unique protease recognition sequences, situated between each of the individual elements that must be separated by the native HIV-1 protease during complete Gag-Pol processing (**SI Fig. 2d**)^46,47^. While these sequences all share some common traits, they are known to be cleaved at varying efficiencies, which ensures that protease processing occurs in the correct order in the native context. In v1 and v2DARTs, the protease-cleavable linker contains the recognition sequence between the matrix and capsid proteins (MA/CA) in endogenous HIV-1. However, across a large set of native and mutated protease cleavage sites, several variants were found to be cut at higher rates than the MA/CA sequence, making them potentially suitable for optimizing DARTs.

We replaced the recognition site in v2DARTs’ protease-cleavable linker with codon-optimized versions of the top three hits from this kinetic screen (**Fig. 3d**)^46^. We then produced v2DARTs and three equivalent formulations with modified linkers, each containing a B2M-targeting sgRNA, and applied them to unconjugated or pre-stained Jurkat cells. We included v2DARTs with a non-targeting sgRNA as a negative control. Among all linker variants, replacing the MA/CA site with the wild-type HIV-1 spacer peptide 1 and nucleocapsid protein (SP1/NC) sequence produced the greatest efficacy, generating two times more knockout at the experimental dose (**Fig. 3e, left**). Notably, all tested variants showed negligible gene editing when targeting antibodies were absent, demonstrating that the unconjugated DARTs remained transduction-incompetent.

To evaluate the mechanism underlying the high potency of the SP1/NC architecture, we measured proteolytic activity by Western blotting. We initially hypothesized that the improved editing efficiency was driven by a simple acceleration of Cas9 cargo liberation; given the reportedly higher cleavage rates of the optimized linkers, we expected the ratio of Gag-Cas9 (220 kDa) to Cas9 alone (160 kDa) to decrease in the new formulations, which we assumed to be desirable. However, our results revealed that this ratio remained relatively invariant between baseline v2DARTs and the formulations with alternative protease-cleavable linkers (**Fig. 3f**). This finding indicated that the functional improvement seen in the SP1/NC particle was not due to increased Cas9 release.

Unexpectedly, visualization of the structural Gag polyprotein (p55) showed that the SP1/NC sequence did significantly impact overall proteolytic processing of DARTs. Staining for p24, the CA domain of the Gag polyprotein, is a useful method for determining the maturation state of a lentiviral particle. The presence of p24 indicates that the structural Gag protein has been fully cleaved by the viral protease; conversely, the presence of p55 or intermediate species (between 55 and 24 kDa) indicates no or partial processing of Gag, respectively. Standard HIV-1 presents with p24 as the predominant species^48^. Due to their lack of maturation (**Fig. 3c**), v2DARTs contain mostly p55, while also exhibiting clear p24 and intermediate bands; a similar profile is seen in R01 RTH/IN and S09 SP1/NC formulations. In contrast, our high-potency SP1/NC DARTs exhibited attenuated proteolytic processing, characterized by reduced levels of both mature p24 and partially cleaved intermediates compared to other formulations; this finding was consistently observed across experiments (**Fig. 3f**, **SI Fig. 2e**).

This inverse correlation between editing rate and protease-mediated maturity was unexpected. The data suggests that the SP1/NC linker produces a particle with an altered internal structure, with significantly less full and partial proteolytic processing than baseline v2DARTs. We postulate that the altered proteolytic kinetics create an inner membrane structural state that facilitates cytosolic payload packaging and release, thereby improving functional cargo delivery to target cells. These findings could inform more rational VLP design in the future; it is currently unknown whether the mature, immature, or undefined state (**Fig. 3c**) yields the most potent particles. However, more work is required to isolate the complete rationale for v3DARTs’ improved function and apply this information to VLP engineering.

Finally, to maximize the probability that the liberated cargo reaches the genome, we optimized the nuclear import efficiency. While VLP packaging requires a balance of nuclear export signals (NES) and localization signals (NLS), editing post-entry relies on Cas9 nuclear localization^10,11,16,18,19^. We appended two additional SV40 NLSs at the C-terminus of the Gag-Cas9 fusion, for a total of four. This modification modestly enhanced editing across the two best linker variants, indicating that nuclear entry remained a relevant rate-limiting step (**Fig. 3e, right**). As such, we designated the final optimized architecture, containing the SP1/NC protease cleavage site and four SV40 NLSs, as v3DARTs (**Fig. 3g**).

Notably, repeating the quantification of packaging fidelity, we identified that v3DARTs incorporated 28% fewer Cas9 molecules than v2DARTs (**Fig. 3h**). This finding supports the earlier conclusion that the internal processing of v3DARTs differs from that of v2DARTs, as the functional improvement of the former is not simply due to the delivery of more Cas9 proteins.

To robustly compare all DART VLP versions, a characteristic dose curve of v3DARTs was produced in the same manner as with the other iterations (**Fig. 3i**). v3DARTs outperformed versions 1 and 2 across all tested antibody concentrations (**Fig. 3j-l**) and retained a lack of activity in the absence of targeting antibody (**SI Fig. 2f**). While v3DARTs produced a higher maximum knock out than v1DARTs, they saturated at approximately the same editing rate as v2DARTs. However, version 3 yielded a significantly lower fitted EC50 than version 2 across all antibody conditions. These results indicate that v3DARTs are more potent than v2DARTs but ultimately achieve the same efficacy at saturation.

### Quantitative validation of particle homogeneity and assembly fidelity

We next performed physicochemical characterization to assess DARTs’ batch reproducibility, payload encapsulation, and surface-ligand saturation: all crucial parameters in the manufacturing of targeted delivery vehicles.

We used the Exodus ultrafiltration system to isolate and concentrate intact DARTs, followed by nanoparticle tracking analysis (NTA) to define their physical parameters. DARTs exhibited a highly monodisperse population, with a mean diameter of 118.3 nm and consistent yields (∼3.11 × 10^14^ particles/mL post-concentration; **Fig. 4a**), demonstrating that the addition of generic surface binders did not disrupt particle assembly or stability.

**Figure 4:**
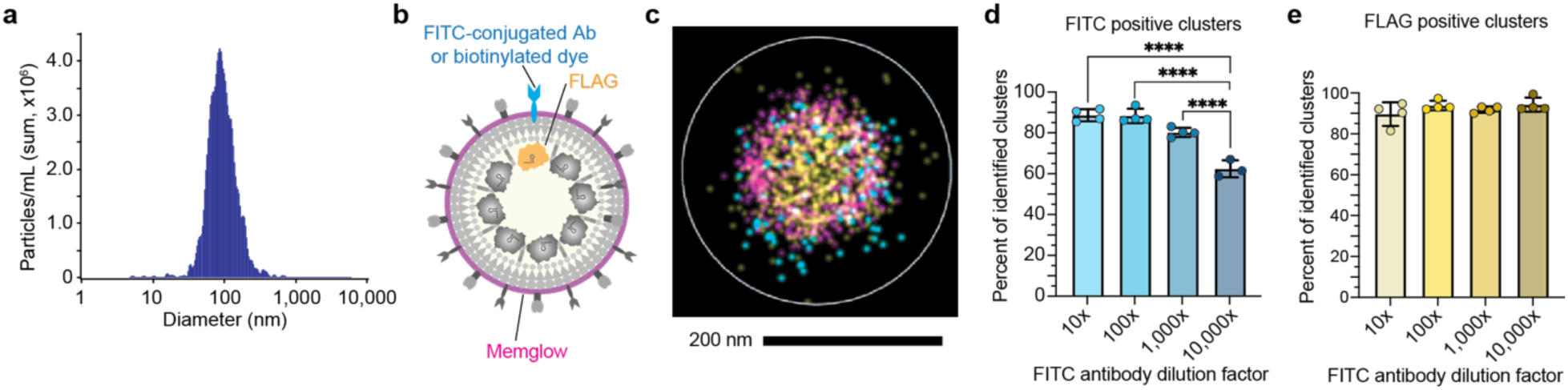
DARTs efficiently conjugate with antibody targeters and encapsulate Cas9 RNP at high rates. **a,** Representative histogram of DARTs diameter, as measured by Nanoparticle Tracking Analysis after concentration via EXODUS. **b,** Diagram of dSTORM imaging strategy for DARTs. Memglow (purple) stains the outer membrane, FLAG (yellow) stains Cas9, and FITC-conjugated antibodies or biotinylated dyes (blue) stain surface-bound generic binders. **c,** Representative image from dSTORM imaging of DARTs. FLAG can be identified within the particle, while Memglow and generic binders are visible on the external surface. Scale bar = 200 nm. **d,e** αCD184-FITC was serially diluted and used to stain αFITC DARTs prior to dSTORM imaging. Percent of identified clusters that were FITC positive (**d**) and FLAG positive (**e**) are plotted. Clusters are called via software intrinsic to the imaging machinery; to curate the set of “identified clusters” used for analysis herein, particles positive for one fluorophore only were excluded to avoid counting antibody and dye aggregates (see **Methods**). All groups were compared to one another via ordinary one-way ANOVA; for all indicated significant differences, P < 0.0001. Data was collected as one lane per sample and averaged across four fields of view per lane. Data are all mean ± s.d. One outlier was identified from the 10,000x group in **d**, as per ROUT test with a false discovery rate of 5%, and subsequently removed from analysis. Antibody clone used for imaging: αCD184, 2B11.

We next employed direct stochastic optical reconstruction microscopy (dSTORM) to characterize the supramolecular architecture of individual DART particles. The outer membranes of the VLPs were stained with MemGlow700, Cas9 was visualized via FLAG staining, and FITC-conjugated antibodies or biotinylated dyes were used to characterize the generic capture binders (**Fig. 4b**). As anticipated, spherical particles were identified with internal Cas9 and surface-tethered mSA or αFITC scFv (**Fig. 4c**, **SI Fig. 2g**). Across imaged particles, FLAG and surface binder fluorophore counts ranged from 138 to 708 and 64 to 183 per particle, respectively (though these numbers do not necessarily correlate 1:1 with the number of actual Cas9 and generic binder molecules).

Our single-particle analysis revealed that the external surface of DARTs is highly efficient at antibody capture. Over a three-log dynamic range of antibody concentrations (1:10 to 1:1,000 dilution), >80% of all “identified particles” – defined as clusters positive for two or more of our fluorophores, to exclude nonspecific signal and antibody or dye aggregates – successfully conjugated with the targeting antibody (**Fig. 4d**, **SI Fig. 2h**). Conjugation rates only significantly decreased at a 1:10,000 dilution, with approximately 60% of DARTs capturing antibody in this condition. These data speak to the potential for DARTs to retain bound antibody even in dilute regimes, as may be encountered with certain *in vivo* injection routes.

We also assessed the cargo encapsulation fidelity, i.e., the probability that a VLP carries the Cas9 editor. Across all tested conditions, >90% of the identified particles contained the Cas9 payload (**Fig. 4e**, **SI Fig. 2h**). This high degree of cargo packaging indicates that the DART platform does not disrupt VLP assembly.

### Programming specific and cooperative entry in primary human T cells

After fully optimizing and characterizing DARTs and demonstrating their efficacy in cell lines, we sought to program diverse targeting logic into more complex systems. Primary human T cells are a clinically-relevant cell type that naturally exists as a heterogeneous mixture of CD4^+^ and CD8^+^ subsets^27^, which is an ideal paradigm for evaluating the specificity and cooperativity of DART-mediated targeting.

Given the modularity of the DART technology, we identified our ability to rapidly probe whether simultaneous engagement of multiple receptors could drive cooperative nanoparticle entry. Leveraging the strengths of our platform, we applied αFITC v3DARTs to primary human T cells using both single antibodies and binary combinations (e.g., αCD4 + αCD7). While single-antigen targeting could be effective, combinatorial targeting revealed a strong synergistic effect (**Fig. 5a**). For example, coupling a subset-specific binder (e.g., αCD4 or αCD8) with a pan-T cell binder (e.g., αCD7 or αCD3) consistently enhanced editing efficiencies beyond the additive potential of either antibody alone (**Fig. 5a**).

**Figure 5:**
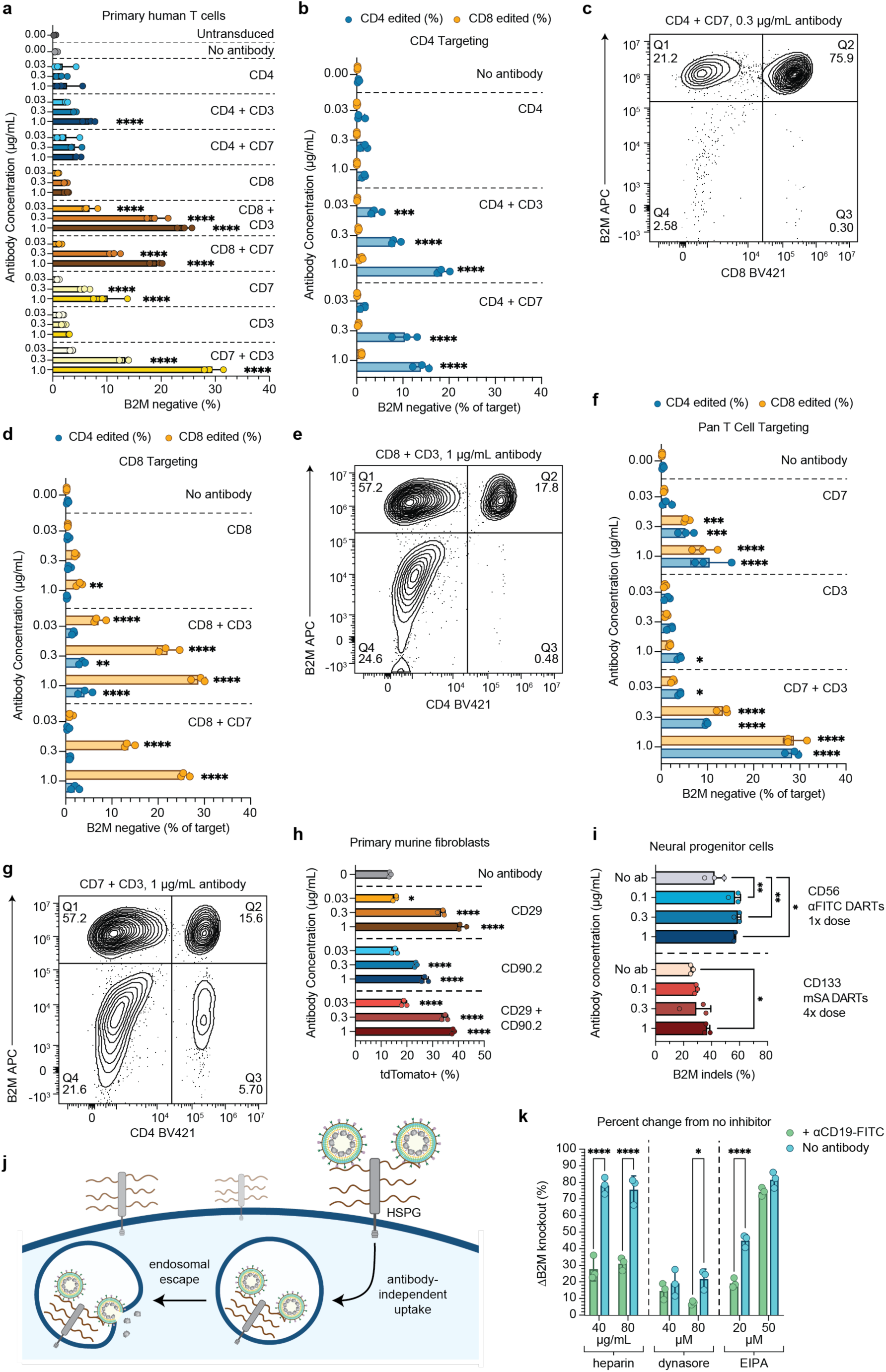
DARTs exhibit high specificity and can transduce complex cell types. **a,** A 4x dose of αFITC v3DARTs carrying sgB2M was targeted towards one primary human T cell donor via an antibody panel. B2M knockout was assayed via flow cytometry 6 days post-transduction. Values are representative of all T cells, irrespective of subtype. **b,d,f** Analysis of **a**. Graphs display the percentage of CD4^+^ and CD8^+^ T cells with B2M knockout when targeting antibodies include αCD4 (**b**), αCD8 (**d**), or only pan T cell markers (**f**). Each group is compared to the no antibody condition of the same T cell subtype via two-way ANOVA. **c,e,g**, Raw flow plots of select conditions from **b**, **d**, and **f**, respectively. **h,** A 1x dose of αFITC v3DARTs carrying sgLoxP were used to transduce primary fibroblasts derived from Ai9 mice. tdTomato signal was quantified by flow cytometry 4 days post-transduction. Media was aspirated and replaced 2 hours post-transduction. **i,** A 1x dose of αFITC and 4x dose of mSA v3DARTs were used to knockout B2M in *NGN2*-derived neural progenitor cells. v3DARTs were directed with αCD56-FITC and αCD133-biotin. Sanger sequencing and ICE analysis were used to quantify indels. DNA extraction was 4 days post-transduction. One outlier was identified via ROUT test (false discovery rate of 5%) and removed. **j,** Heparan sulfate proteoglycan (HSPG)-mediated background transduction by naïve DARTs. **k,** αFITC v3DARTs carrying sgB2M were targeted to CD19-expressing HEK cells with or without transduction inhibitors. HEK cells were unmodified or pre-treated with αCD19-FITC. For each inhibitor concentration and antibody treatment, percent change in B2M knockout from the no inhibitor condition is plotted. Two-way ANOVA was used to compare +/− antibody conditions within the same inhibitor treatment. For all data, n=3 technical replicates per condition. Unless otherwise specified, conditions were compared to the no antibody control via two-way ANOVA. Data are all mean ± s.d. For all plots: *, P<0.05; **, P<0.01; ***, P<0.001; ****, P<0.0001. Antibody clones used for transduction: αCD4, SK3; αCD3, HIT3a; αCD7, CD7-6B7; αCD8, RPA-T8; αCD29, HMβ1-1; αCD90.2, 30-H12; αCD56, 5.1H11; αCD133, AC133; αCD19; 4G7. Antibody clones used for staining: αB2M, 2M2; αCD4, SK3; αCD8, RPA-T8. Abbreviations: B2M, β-2 microglobulin; HSPG, heparan sulfate proteoglycan.

Crucially, this cooperativity was still governed by strong cell-type specificity. When an αCD4 antibody was used, significant editing occurred only in CD4^+^ populations, even when combined with an αCD3 or αCD7 antibody that should bind all T cells (**Fig. 5b,c**, **SI Fig. 3a**). The same preferential targeting largely held for CD8^+^ cells, with no delivery to CD4^+^ populations evident in the αCD7 combination conditions; the addition of αCD3 did produce low rates of off-target editing, though the preference remained for CD8^+^ cells in these samples (**Fig. 5d,e**, **SI Fig. 3b**). When only pan T cell markers were used to program DARTs, roughly equal fractions of CD4^+^ and CD8^+^ cells were edited (**Fig. 5f,g**, **SI Fig. 3c**). These data may suggest that, in certain cases, a highly specific interaction can direct nanoparticle binding while a more general marker facilitates internalization. Such a scenario might occur when the more specific surface receptor endocytoses more slowly, while the general one rapidly turns over; the specific marker directs binding, while the nanoparticle leverages the general marker’s consistent internalization for entry. This hypothesis is in line with literature indicating that different receptors are endocytosed (1) at significantly different rates and (2) in response to different stimuli, with some naturally turning over and some internalizing in a ligand-dependent manner^49–53^.

Whether the observed paradigm is generalizable to broader contexts warrants further study, as it could be leveraged powerfully in many applications. However, it is worth noting that complex antigen interactions such as these would be challenging to identify without a modular system such as the DART platform, which enables simultaneous testing of many antibody combinations. This entire process was repeated with a second blood donor at a second genomic locus, yielding similar results (**SI Fig. 3d-g**).

### Diverse, programmable antigen targeting in adherent cell types

To demonstrate the platform’s cross-species and cross-tissue universality, we also applied v3DARTs to primary murine fibroblasts and human induced pluripotent stem cell (iPSC)-derived neural progenitor cells (NPCs). We isolated the fibroblasts from Ai9 reporter mice, in which Cas9 cleavage at two LoxP sites can excise a stop codon and induce tdTomato expression (however, it is worth noting that tdTomato positivity underreports Cas9 delivery and editing, since only a fraction of edits will actually activate the cassette). Via modular antibody conjugation at various concentrations, we identified CD29 and CD90.2 as functional entry targets, achieving robust reporter expression (**Fig. 5h**). v3DARTs were also capable of knocking out B2M in iPSC-derived NPC populations, a difficult-to-transduce cell type^56,57^; v3DARTs targeted against CD56 or CD133 achieved strong B2M knockout rates of up to 60% in this context (**Fig. 5i**).

When performing gene knockout in adherent cell types, such as fibroblasts and NPCs, it became evident that the background transduction level from naïve DARTs was significantly higher than in suspension cells. We first hypothesized that this finding was attributable to gravitational sedimentation, with non-targeted particles settling onto the cell monolayer and undergoing non-specific, contact-dependent endocytosis. However, this theory was not deeply explanatory. To probe the phenomenon further, we first generated an adherent HEK293T cell line expressing CD19 fused to a GFP reporter, as previously described^10^. We then added αFITC v3DARTs to these cells with and without targeting antibody and in the presence or absence of three small-molecule inhibitors: heparin, which disrupts interactions with heparan sulfate proteoglycans (HSPGs)^58^; dynasore, which inhibits dynamin-1 and −2 to disrupt clathrin-mediated endocytosis^59,60^; and ethylisopropylamiloride (EIPA), which blocks sodium-hydrogen exchangers to disrupt macropinocytosis^61,62^. To isolate the nature of our observed non-specific activity in adherent cells, we hoped to identify inhibitors that specifically impacted DARTs’ antibody-independent activity.

It is known that certain viruses and nanoparticles, such as SARS-Cov-2^63,64^ and VLPs^58^, interact with HSPGs in addition to their more canonical entry receptors to facilitate cell uptake^65^. We therefore hypothesized that DARTs may be operating via a similar mechanism to enter adherent cells in an unguided manner (**Fig. 5j**). Indeed, the addition of heparin and subsequent blocking of HSPG binding significantly inhibited DARTs’ antibody-independent activity, while minimally impacting antibody-directed editing (**Fig. 5k**, **SI Fig. 3h**). These data suggest that binding of HSPGs in adherent cell types is a major contributor to background activity, the knowledge of which can inform strategies for inhibiting undesirable non-specific uptake.

This experiment also showed significant inhibition of antibody-independent (at 20μM) and overall transduction (at 50μM) by EIPA (**Fig. 5k**). While these findings may implicate macropinocytosis in nonspecific uptake of DARTs, they also may be an artifact of EIPA increasing endosome pH, which inhibits mVSV-G-mediated endosomal escape. Further study is required to isolate the exact role of macropinocytosis in DARTs entry.

### DART is agnostic towards VLP scaffold and packaged cargo

While HIV-1-based VLPs are a promising platform, similar technologies based on Moloney Murine Leukemia Virus (MMLV) scaffolds have recently shown even higher efficacy rates^13,21^. Because the DART system should interact only with the outer envelope of its delivery vehicle and interface minimally with its internal components, we rationalized that DART should also be compatible with MMLV-derived particles packaging alternative cargoes.

To test this hypothesis, we appended the DART technology to a highly optimized, MMLV-based VLP carrying an adenine base editor (ABE)^21^; we termed the resulting vehicles base editing DARTs (be-DARTs). Using prior methods, we produced a dose curve in a base-editing Jurkat cell reporter line, in which successful A◊G conversions induce GFP expressio^66^. We found be-DARTs to be highly potent, with successful editing observed in approximately 100% of cells at doses above 20μL (**Fig. 6a**). Minimal antibody-independent activity was seen only at severely saturating doses. Notably, be-DARTs’ efficacy closely mirrored that of the published, standard VSV-G pseudotyped particles.

**Figure 6:**
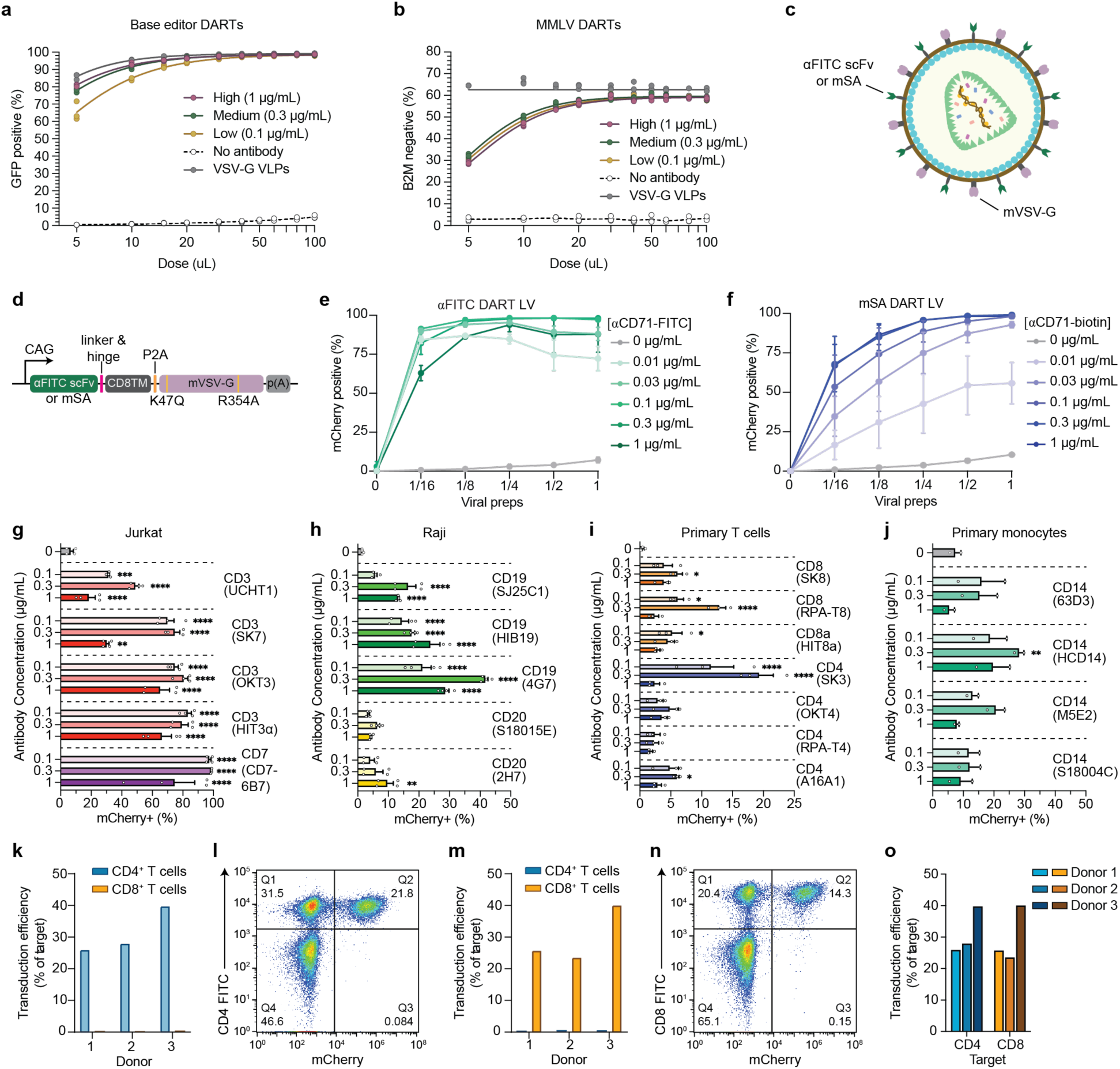
The DART targeting system is compatible with multiple packaged cargoes and alternative viral and non-viral scaffolds. **a,** MMLV-derived base editor DARTs (be-DARTs) were applied to a GFP reporter line to produce a dose curve, as in **Fig. 2c**. be-DARTs were compared to analogous MMLV-derived base editing VLPs pseudotyped with wild-type VSV-G (grey). **b,** A dose curve was produced using Cas9-packaging DARTs with MMLV-based internal components (termed MMLV DARTs), as in **Fig. 2c**. MMLV DARTs were compared to a VSV-G pseudotyped analog. All replicates are plotted and curves were fitted via sigmoidal four-parameter logistic regression. **c,** Schematic of DART LV, which delivers transgenes for integration. **d,** Diagram of generic binder architectures for DART LV. **e,f,** αFITC scFv (**e**) and mSA (**f**) decorated lentiviruses were incubated with FITC-conjugated or biotinylated αCD71 antibodies, respectively, and used to transduce an mCherry expression cassette into Jurkat cells. Total n=6, with two experiments performed in technical triplicate. **g,** Using the full catalog of FITC-conjugated αCD3 and αCD7 antibodies available from BioLegend, a 1x dose of αFITC DART LV was targeted to Jurkat cells. **h,** As in **g**, αCD19 and αCD20 clones were used to transduce Raji cells. **i,** As in **g**, primary human T cells were transduced by DART LV targeted with αCD4 and αCD8 antibodies. All data in **g**-**i** is n=4, with two biological replicates in technical duplicate. **j,** As in **g**, lentivirus-resistant primary monocytes were transduced by DART LV conjugated to αCD14-FITC. Total n=2 technical replicates. **k-n** DART LV was used to transduce an mCherry expression cassette into primary human T cells from three different blood donors. DART LV was targeted either with αCD4-FITC (**k,l**) or αCD8-FITC (**m,n**). **o,** Side-by-side comparison of **k** and **m**. All data in **k**-**o** is n=3 biological replicates. A 1x dose of αFITC DART LV was used for all conditions in **g**-**o**. For **e**-**o**, mCherry positivity was assessed via flow cytometry 5 days post-transduction. All plotted data with error bars is mean ± s.e.m. Antibody clones used for transduction and staining, unless otherwise stated: αCD3, HIT3a; αCD71, OKT9; αCD4, SK3 (**i-m**); αCD8, RPA-T8 (**i-m**). Abbreviations: αFITC scFv, anti-FITC single-chain variable fragment; mSA, monomeric streptavidin; CD8TM, CD8α transmembrane domain; p(A), poly-A tail.

To contribute to head-to-head benchmarking of HIV-1- and MMLV-based platforms, we also attached the DART technology to optimized MMLV VLPs – i.e., the published v5 eVLPs^21^ – packaging Cas9 for genome editing; we simply named these MMLV DARTs. These, too, worked as anticipated, achieving saturation at a 30μL dose and exhibiting no antibody-independent activity (**Fig. 6b**). Notably, while MMLV DARTs underperformed their VSV-G pseudotyped counterpart at low doses, they achieved near parity with higher amounts. Additionally, MMLV DARTs outperformed HIV-1-derived v3DARTs across all tested doses and antibody concentrations, with the two formulations exhibiting similar off-target profiles (**SI Fig. 3i**).

### DART can be applied to other virus-based systems

To further demonstrate the universality of the DART technology, we next transferred its targeting logic to an integrative lentiviral vector system. Given the prior results, we hypothesized that the DART platform is internally agnostic, capable of routing any membrane-enveloped vehicle regardless of its structure or payload. Accordingly, we generated DART lentivirus (DART LV) with generic capture binders (mSA or αFITC scFv) encoded upstream of mVSV-G, connected by a P2A self-cleaving peptide (**Fig. 6c,d**).

To characterize DART LV, we again established a dose curve in Jurkat cells over various antibody concentrations. In this instance, we chose to target CD71; also known as transferrin receptor 1 (TfR1), it is a ubiquitous transmembrane glycoprotein essential for iron uptake, which is often leveraged for cellular entry via receptor-mediated endocytosis^67^.

Both αFITC and mSA DART LV could transduce up to 100% of cells, exhibiting clear dose responsiveness to viral and antibody inputs (**Fig. 6e,f**). In the absence of a targeting antibody, DART LV performed minimal transduction, demonstrating the same high specificity as our VLPs. Notably, by testing DART LV across a larger antibody dose regime, we obtained further support for the Hook effect observed in our prior dose curves; with αFITC DART LV, one can observe transduction inhibition at the highest antibody concentration (**Fig. 6e**). As such, this data helps establish important stoichiometric parameters governing optimal DART assembly.

One powerful feature of DART LV is its ability to evaluate the viral entry capacity of antibody clones targeting different epitopes on the same receptor. Conventional vector engineering treats an antigen (e.g., CD3) as a whole, single target. However, by screening a panel of antibody clones against identical cell surface markers – including CD3 (Jurkats, primary human T cells), CD19 and CD20 (Raji cells), CD8 and CD4 (primary human T cells), and CD14 (primary human monocytes) – DART LV revealed differences in internalization efficiencies between antibody clones targeting the same protein (**Fig. 6g-j**). While some of these transduction discrepancies can be explained by differences in antibody affinity, K_D_ values are not generally available for every commercial antibody or binder. As such, DART LV can function as an upstream “discovery engine” to empirically identify the most transduction-competent antibody clones, providing a more defined blueprint for future design of membrane-integrated scFvs or nanobodies.

Another crucial aspect of DART LV is its ability to transduce cell types resistant to standard VSV-G lentiviral pseudotypes, such as primary human monocytes^68^. We leveraged our platform’s programmable tropism to overcome lineage-specific engineering barriers, successfully transducing primary human monocytes by re-routing the virus through the CD14 receptor (**Fig. 6j**).

Finally, we also demonstrated that DART LV retains the high cell-type specificity characteristic of our platform. Primary human T cells from three donors were transduced with DART LV conjugated to either αCD4-FITC or αCD8-FITC; CD4-targeted DART LV transduced CD4^+^ populations exclusively (**Fig. 6k,l**), while CD8-targeted DART LV transduced CD8^+^ T cells exclusively (**Fig. 6m,n**). In both cases, DART LV was highly efficient, with transduction efficiencies reaching 40% of on-target cells in the CD4^+^ and CD8^+^ populations (**Fig. 6o**). Altogether, these results confirm that the DART interface is modular and internally agnostic, and that it expands accessible therapeutic reach to difficult-to-target cell lineages.

### DART-mediated antibody screening reveals CD56 as a functional receptor for neural progenitor targeting in the fetal brain

One major hurdle hampering delivery is the introduction of gene therapies into neural populations *in vivo*^1,3,5^. Though VLP formulations have been tested in the murine and non-human primate brain, they have either failed to transduce neurons or to specifically target them^11,14,18^. Furthermore, efforts to engineer VLPs for neuron-specific targeting, such as pseudotyping with neurotrophic viral glycoproteins^14^, are lengthy, complex, and often inefficient. This challenge is exacerbated by the fact that targeting tropism within the central nervous system (CNS) is governed by complex tissue architectures and transport dynamics that cannot be easily modeled in culture.

To grapple with this persistent barrier, we deployed DARTs as an *in vivo* antigen discovery platform, seeking to identify optimal entry receptors on neural progenitor cells (NPCs) in the developing brain. We chose to administer them to mouse fetuses *in utero* at embryonic day 15.5 (E15.5), as this is a critical window for murine cortical neurogenesis^69,70^.

We first identified which cell surface markers were characteristic of NPCs at the E15.5 developmental stage and assessed which of our candidate commercial antibodies were strong and efficient binders. C-sections were performed to liberate E15.5 mouse pups, and their brain tissue was digested into a single-cell suspension and stained with prospective targeting antibodies. Candidate antigens included CD56, CD133, CD166, and CD184. While the literature suggests that all of these candidates are present on E15.5 NPCs, only one αCD56-FITC clone produced a distinct, brightly-stained population separated from the background (**Fig. 7a**); the other antibodies generated less distinct staining patterns or a broadly shifted background, making them less suitable candidates for *in vivo* targeting (**SI Fig. 4b-k**). Crucially, no public scFv sequences exist for murine CD56, rendering this receptor inaccessible to VLPs containing hard-coded, membrane-bound targeters without an extensive scFv discovery procedure.

**Figure 7:**
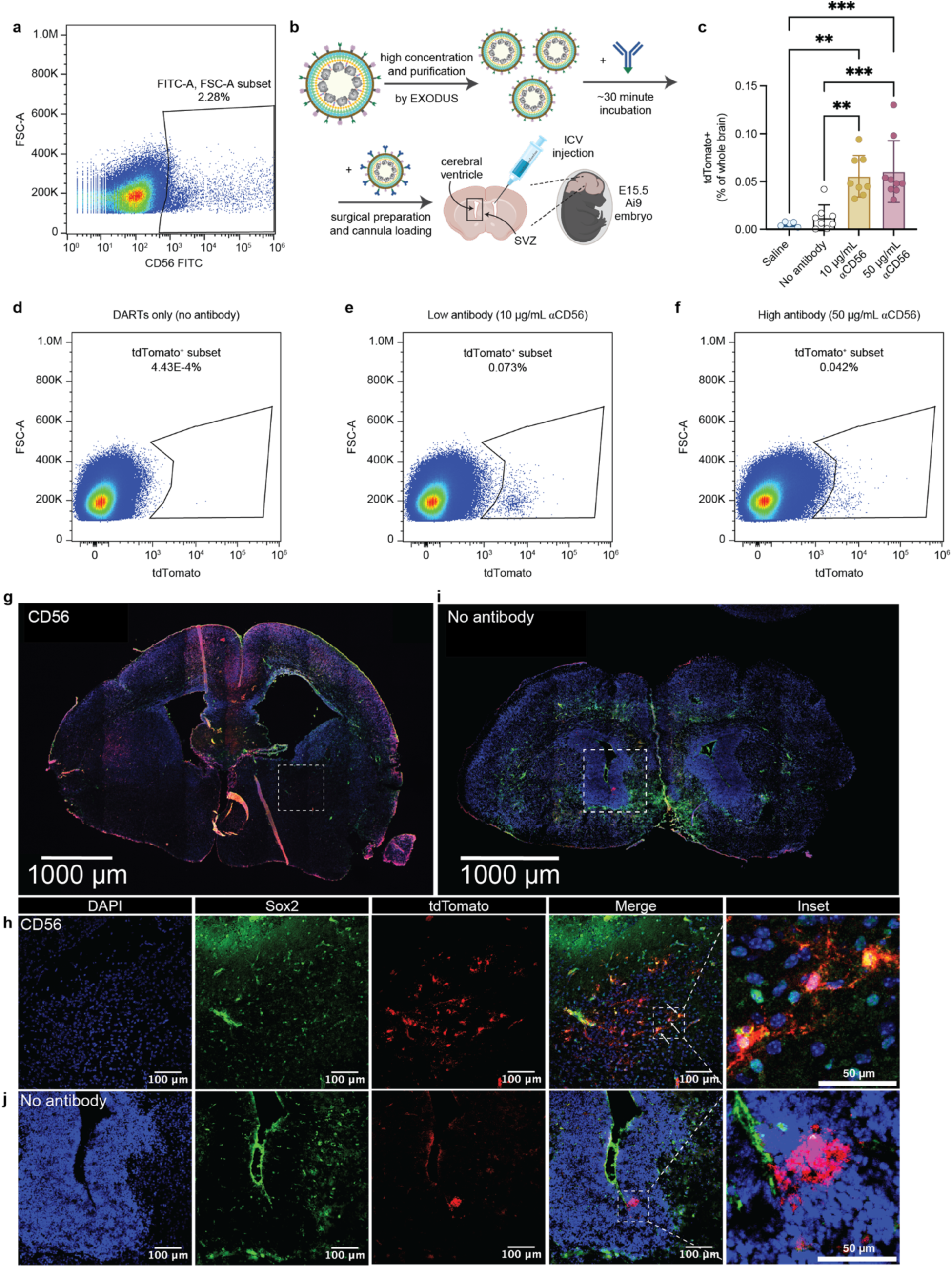
DARTs can identify target antigens for transduction of neural cell populations and perform gene editing *in vivo*. **a,** E15.5 Ai9 fetal mouse brains were harvested, digested, and stained with a panel of FITC-conjugated antibodies. The αCD56-FITC antibody shown was progressed for *in vivo* use. **b,** Schematic of the workflow for *in vivo* experimentation. DARTs were highly concentrated by EXODUS and incubated with targeting antibody at low (10µg/mL) or high (50µg/mL) concentration while surgical preparation was performed. Low and high concentrations were chosen by scaling the antibody dose linearly with VLP concentration increases from *in vitro* to *in vivo* implementation. DARTs were then injected into fetuses *in utero* via intracerebroventricular injection targeting the subventricular zone (SVZ). **c,** Whole-brain flow cytometry analysis of E19.5 fetal mouse brains 4 days post-injection. Fetuses were treated with saline, naïve DARTs, or DARTs conjugated with 10 or 50µg/mL αCD56-FITC. n=5-9 biological replicates per group. Injections were performed in 3 dams across 3 independent experiments, with fetuses from each litter assigned to multiple treatment groups to minimize confounding between treatment and litter. Litters were not separated statistically. **d-f,** Raw flow plots from representative no antibody (**d**), low antibody (**e**), and high antibody (**f**) samples plotted in **c**. An additional set of plots can be found at **SI Fig. 5a-c**. **g,** Preserved brain section from one Ai9 mouse fetus treated with αFITC v3DARTs targeted by αCD56-FITC at 50 µg/mL. **h,** Higher magnification of the region in **g** contained in the white box. Arrows indicate highlighted tdTomato^+^/Sox2^+^ cells depicted within the inset. **i,** Preserved brain section from an Ai9 mouse fetus treated with naïve αFITC v3DARTs not conjugated to an antibody. **j,** Higher magnification of the region in **f** contained in the white box. (**g,i**) Scale bars = 1 mm. (**h,j**) Scale bars = 100 µm, inset scale bars = 50 µm. Antibody clone used for transduction: αCD56, polyclonal. Antibody clones used for staining: αSox2, L1D6A2; αtdTomato, polyclonal. Abbreviations: SVZ, subventricular zone; ICV injection, intracerebroventricular injection.

Having identified CD56 as a candidate antigen, we injected naïve and CD56-programmed DARTs directly into E15.5 fetuses via the intracerebroventricular (ICV) route (**Fig. 7b**). DARTs packaging an sgRNA targeting LoxP were injected into pups from the Ai9 mouse line, wherein Cas9 editing induces tdTomato expression in a fraction of cases (**SI Fig. 4a**)^54,55^. C-section was performed at E19.5, treated pups were dissected, and brain tissue was either digested for flow cytometry or stained for microscopy. E19.5 whole-brain analysis via flow cytometry revealed pronounced editing in mouse pups treated with CD56-directed DARTs, but not naïve DARTs or the saline control (**Fig. 7c-f**, **SI Fig. 5a-c**). To directly analyze their target-cell specificity, brain sections were stained for neuroepithelial stem cell protein (Nestin) and SRY-box transcription factor 2 (Sox2), proteins which characterize NPCs at the analytical time point^71–74^.

Crucially, image-based analysis of brain sections treated with CD56-targeted DARTs (**Fig. 7g,h**, **SI Fig. 5d-g**) versus DARTs alone (**Fig. 7i,j**) revealed that only the antibody-conjugated particles edited NPC populations. In a representative brain section from a fetus treated with αCD56-conjugated DARTs (at 50 μg/mL antibody), the targeted nanoparticles exhibited distinct tropism-directed editing patterns in two regions adjacent to the cerebral ventricles. Within these two ventricle-proximal regions (**Fig. 7h**, **SI Fig. 5e**), computational analysis revealed target-cell specificity of 93% and 83%, with 11% and 16% of all Sox2^+^ cells being edited, respectively (**SI Fig. 6a,b**).

In contrast, one brain section from a fetus treated with naïve DARTs (**Fig. 7i**) showed only one small, localized patch of editing near the ventricles and toward the back of the skull (**Fig. 7j**). While naïve DARTs did nonspecifically transduce this small pocket of cells, only 20% of cells within the region of discernable editing were within our “on-target” Sox2^+^ population (**SI Fig. 6c**). As such, while both CD56-targeted and naïve DARTs did edit cells relatively close to injection site, our local measurements indicate that only CD56-directed DARTs showed high tropism for NPCs.

Although the low absolute transduction efficiency (∼11-16% of the Sox2^+^ population in select regions near the subventricular zone (SVZ); **SI Fig. 6a,b**) reflects the inherent physical barriers of *in vivo* nanoparticle diffusion in the brain, our results validate CD56 as a functional antigen for cell type-specific delivery to NPCs. More broadly, our findings demonstrate DARTs’ utility as a rapid prototyping tool. We successfully screened a panel of antigens, identified a receptor with no available genetically encoded binders, and validated its tropism *in vivo* without cloning a single vector for viral pseudotyping. Importantly, because no murine CD56 binders currently exist, the use of the DART system in this context is not only a convenient and rapid method for researchers to identify the receptor’s *in vivo* targeting potential but may also be one of the only pragmatic ways to do so. This example illustrates firsthand how DART technology expands the existing space of targetable cell surface antigens.

## Discussion

The engineering of programmable delivery vehicles is currently constrained by a fundamental, structural bottleneck: the direct coupling between a particle and its targeting entity. Popular tropism retargeting strategies require the genetic hard-coding of fusogens or ligands into a vector’s envelope^10–15,22–33^, forcing a laborious design-build-test cycle that limits exploration of the viral entry potential of the cell-surface proteome. While prior strategies exist to combat this issue via antibody-based targeting, they lack bio-orthogonality or the ability to interface with large pools of commercial antibodies without prerequisite chemical modifications^30^.

We have instead developed the DART platform for rapid empirical assessment of receptors enabling cell type-specific delivery. DART transforms nanoparticle tropism engineering from a bespoke cloning or “guess and check” process into a modular, integrated workflow. By establishing a universal interface for cheap, off-the-shelf antibodies, the DART technology allows users to rapidly probe any surface antigen *in vitro* and *in vivo* to identify cell type-specific targeting parameters without the prerequisite of ligand discovery, modification, or optimization.

A critical insight from our study, which our technological platform also addresses, is that receptor interaction does not guarantee viral entry; this phenomenon may even be exploitable in dual antigen-targeting contexts (**Fig. 5**). It has been noted by others that certain binders do not well facilitate viral uptake, though it remains unclear whether this is a feature of the targeted antigen’s internalization rate, the binder, both, or other factors^10,49,75,76^. As such, while transcriptomics can identify highly expressed surface markers, it cannot predict which receptors or which epitopes on a receptor will mediate productive endocytosis. This fact highlights the necessity of modular, empirical assessment strategies such as DART.

With antigen interaction not directly correlating with viral entry, there is no guarantee that any of the sparsely available scFv or nanobody sequences will be suitable for transduction. Our data highlights this unpredictability: within the CD3 antigen alone, different antibody clones yielded vastly different transduction efficiencies (**Fig. 6g**), and screening revealed CD19 as a functional entry point for Raji cells where CD20 failed (**Fig. 6h**). The result is that users aiming for cell type-specificity are forced to adhere to common targets with many available binder sequences or well-validated delivery precedents, thus limiting the overall scope of targeted delivery approaches.

The DART system therefore bridges the gap between “omics” data and functional delivery, serving as a high-throughput method to map the entry logic of various target tissues and cell types. Furthermore, any identified antigen targets should not be limited to VLP or lentiviral technologies; scFv-based delivery has also been applied to LNPs, another popular delivery system^43,77–79^.

The modularity of the DART platform also unlocks sophisticated combinatorial targeting capabilities. We demonstrate that the surface attachment of multiplexed antibodies can drive synergistic entry into T-cell subsets without loss of specificity (**Fig. 5**). This feature suggests that future delivery systems can and should move beyond single-antigen targeting toward logic-gated entry, mirroring the complexity of physiological cell-cell interactions.

Our optimization of DARTs’ internal structure also supports key biophysical principles for VLP engineering that have been identified in other works^10,11,18,21^. We demonstrate that increasing cargo density and tuning the kinetics of proteolytic processing are critical for maximizing potency (**Fig. 2-3**). While questions remain regarding the optimal maturation status of Cas9-packaging HIV-1-based VLPs, our data support others’ findings that complete virion processing may not be a prerequisite for effective delivery, provided that the RNP payload can be sufficiently liberated from the structural Gag lattice. However, if future work suggests that greater maturation would be beneficial for alternative VLP-mediated delivery, such as when packaging epigenetic editing components, more work is required to determine how to boost proteolytic processing.

Finally, the DART platform addresses the urgent need for robust, practical *in vivo* tools for exploration and validation of cell type-specific delivery. By rapidly screening antigens to target neurons in the fetal brain, we achieved highly specific editing of neural progenitors, demonstrating that surface-based targeting logic can direct biodistribution even in challenging tissues.

While DARTs were introduced into the circulating cerebrospinal fluid (CSF) in this work, a key question remains how they will perform in even more dilutive environments, e.g., after systemic intravenous injection. Evidence herein suggests that non-covalent attachment of antibodies with enveloped delivery vehicles is robust; dSTORM imaging demonstrates that antibodies introduced at low concentration remained tethered to DARTs, even after several washes (**Fig. 4d**). Regardless, further validation of DARTs in diverse *in vivo* contexts is a worthwhile trajectory. If one speculates that antibody detachment poses a severe challenge, they may use the DART system to validate targets *in vitro*, then switch to a similar genetically encoded or identical covalently attached surface-bound targeting moiety for *in vivo* administration.

While DARTs are demonstrably suitable for initial *in vivo* target antigen discovery, they may not be a relevant tool for direct clinical application. While the *in vivo* stability of the antibody linkage is clearly robust enough to mediate cell type-specific delivery (**Fig. 7**), the administration of free antibody is likely to induce an immune response in the host, which may exacerbate a response to immunogenic VLP components^80,81^. For safety purposes, we envision the translational role of the DART platform to be within the target identification stage; it can be employed to identify optimal cell surface antigens mediating highly specific delivery *in vitro* in human cells and *in vivo* in animal models. For subsequent therapeutic vectors, the generic capture binder can be supplanted with a membrane-integrated targeter to eliminate the hazard of detached antibodies, or VLPs can be replaced entirely with an alternative delivery vehicle of interest.

While perhaps not a directly patient-applicable tool, the key contribution of the DART technology is its ability to rapidly and empirically identify transduction-competent cell surface antigens. DART provides a practical platform to test readily-available candidate antibodies and their combinations without rebuilding a nanoparticle for each new target, helping to identify cell type-specific delivery strategies for further development. As the field advances towards safer therapeutic strategies, such as precision epigenetic and base editing, DARTs provide a plug-and-play scaffold to rapidly prototype any delivery vehicle – elevating the sophistication of our targeting logic towards parity with the precision of our gene editor payloads.

## Methods

### Plasmid construction

The VSV-G plasmid (Addgene 8454) was a gift from Bob Weinberg. The mutated VSV-G plasmid (Addgene 201913; referred to as mVSV-G), the scFv entry plasmid (Addgene 201913), the U6-reci Gag-Cas9 plasmid (Addgene 201914; referred to as the Gag-Cas9 sgRNA expression vector), the U6-reci Gag-pol plasmid (Addgene 201915; referred to as the Gag-Pol sgRNA expression vector), and the CD19-EGFP lentiviral transfer plasmid (Addgene 201919) were gifts from Jennifer Doudna. Plasmids psPAX2 (Addgene 12260) and pMD2.G (Addgene 12259) were gifts from Didier Trono. The v5 eVLP MMLV gag-pro-pol plasmid (Addgene 228466), the v5 eVLP adenine base editor plasmid (Addgene 228467), and the v5 eVLP Cas9 plasmid (Addgene 228468) were gifts from David Liu. The pEF-XMAS-1xStop plasmid was a gift from Xiao Wang (Addgene 164411).

Oligos encoding the sgRNA spacers were phosphorylated, annealed, and ligated into BsmBI-digested Gag-Cas9 and Gag-Pol expression vectors. mSA^38^ and anti-FITC scFv^36^ were reported in the literature and synthesized by Twist Bioscience (South San Francisco, USA); generic binder plasmids were then generated by in-fusion cloning (Takara Bio) with the scFv entry plasmid opened by BsmBI and EcoRI or NheI and EcoRI. scFv entry plasmid derivatives, Gag-Cas9 plasmid derivatives, and Gag-Pol plasmid derivatives were generated from Addgene 201912, 201914, and 201915, respectively, using in-fusion cloning (Takara Bio). The new protease cut sites and additional NLS sequences were introduced with overlapping PCRs. The base editing reporter plasmid (pSLQ17937) was generated from Addgene 164411 as described previously^66^.

For DART LV, pMD2.G was first mutated via PCR to introduce a K47Q and R354A mutation in VSV-G. Then mSA or the αFITC scFv, a linker, and a CD8 transmembrane domain were introduced upstream of mVSV-G and separated by a P2A tag, creating pSLQ11875 and pSLQ11876.

### Culture of cancer cell lines

HEK293T and Lenti-X cells (Clontech) were cultured in DMEM + GlutaMAX (ThermoFisher) supplemented with 10% fetal bovine serum (Alstem) and 100U/mL of penicillin and streptomycin (Life Technologies). Jurkat cells and Raji cells (ATCC) were cultured in RPMI 1640 + HEPES + L-glutamine (ThermoFisher) supplemented with 10% fetal bovine serum (Alstem) and 100U/mL of penicillin and streptomycin (Life Technologies). Cells were maintained at 37°C and 5% CO_2_ and passaged using standard cell culture techniques.

### Cell line generation

To produce lentivirus for cell line generation, Lenti-X cells were seeded in 6-well plates (Corning) at a density of 1.2e6 cells per dish (Corning). After approximately 16-18 hours, cells were transfected using TransIT-LT1 reagent (Mirus Bio) according to the manufacturer’s instructions. Lentivirus was produced by transfecting a plasmid mixture of psPAX2 (1.3µg), pMD2.G (555ng), and the appropriate transfer plasmid (1.8µg) in each well. 6 hours post-transfection, media was changed to DMEM + GlutaMAX (ThermoFisher) supplemented with 10% fetal bovine serum (Alstem), 100U/mL of penicillin and streptomycin (Life Technologies), and 1x ViralBoost Reagent (Alstem). 24 hours post-transfection, lentivirus was harvested and filtered through 0.45µm syringe filters (Fisher Scientific). Lentivirus was either used immediately or, if indicated, concentrated overnight using Lenti-X solution (Alstem) according to the manufacturer’s instructions.

CD19-expressing cells were generated by transducing HEK293T cells (3e5 cells per well in a 6-well dish) with 1mL unconcentrated lentivirus containing the CD19-EGFP lentiviral transfer plasmid (Addgene 201919) in 2mL total volume per well. Five days post-transduction, flow cytometry was used to evaluate EGFP expression. To generate a population of ∼100% CD19+, EGFP+ cells, the transduced cells were sorted for EGFP expression using an SH800S cell sorter (Sony Biotechnology). CD19 expression was further confirmed via staining with an APC anti-human CD19 antibody.

The base editing reporter cell line was generated by transducing Jurkat cells (3e5 cells per well in a 6-well dish) with 1x concentrated lentivirus containing the base editing reporter lentiviral transfer plasmid (pSLQ17937) in 2mL total volume per well; the base editing reporter contains a constitutive BFP marker and a GFP cassette that turns on in response to an A◊G base edit^66^. To generate a ∼100% transduced population, the transduced cells were sorted for BFP expression using an SH800S cell sorter (Sony Biotechnology).

### Primary T cell isolation and culture

Buffy coats (Stanford Blood Center) or leukopaks (STEMCELL) derived from anonymous healthy blood donors were purchased under an IRB-exempt protocol. Primary human T cells were isolated using the EasySep Human T cell Isolation kit (STEMCELL) according to the manufacturer’s protocol with Ficoll-Paque PLUS (GE Healthcare) and SepMate-50 tubes as needed. Isolated T cells were immediately cryopreserved at 4-10e6 cells per vial in FBS supplemented with 10% DMSO (Sigma-Aldrich).

Cryopreserved T cells were thawed and activated same-day using Human T-cell activator CD3/CD28 Dynabeads (Gibco) at a 1:1 bead to cell ratio. T cells were cultured in RPMI 1640 + HEPES + L-glutamine (ThermoFisher) supplemented with 10% fetal bovine serum (Alstem), 100U/mL of penicillin and streptomycin (Life Technologies), and 200U/mL of human recombinant IL-2 (STEMCELL). Dynabeads were magnetically removed on day two or three of culture (one or two days post-transduction). Cells were maintained at an overall concentration of 0.5-1e6 cells/mL. Live cell counts were obtained following the manufacturer’s protocol for Trypan Blue exclusion using a Countess 3 automated cell counter (Invitrogen).

### Primary monocyte isolation and culture

CD14^+^ monocytes were isolated by EasySep immunomagnetic positive separation (STEMCELL) per the manufacturer’s instructions and used immediately. Primary monocytes were cultured in RPMI + HEPES + L-glutamine (ThermoFisher) supplemented with 10% FBS (Alstem), 100U/mL of penicillin and streptomycin (Life Technologies), and 10ng/mL of GM-CSF (STEMCELL) at a density of 1e6 cells/mL and were not subcultured before assaying.

### Primary Ai9 fibroblast isolation and culture

Primary dermal fibroblasts were isolated from Ai9 (Gt(ROSA)26Sor) reporter mice following euthanasia. To ensure sterility, hair was removed using a topical depilatory agent (Nair), and the carcass was submerged sequentially in povidone-iodine (Medline) and 70% ethanol for 10 minutes each. Ear pinnae tissue was harvested, washed serially through six wells of 70% ethanol and six wells of sterile PBS, and finely minced into fragments (<1mm³). The tissue was digested in 1.5mL tubes containing 500µL of 1mg/mL collagenase in PBS at 37°C for 45 minutes, with vortexing every 15 minutes to facilitate dissociation. Following digestion, samples were centrifuged at 300xg for 5 minutes, and the pellet was resuspended in a minimal volume of D10 medium (DMEM supplemented with 10% fetal bovine serum (Alstem) and 1% penicillin-streptomycin (Life Technologies)). The tissue suspension was plated using a low-volume explant technique to rely on surface tension for initial adhesion; cultures were maintained at 37°C with 5% CO₂, and additional medium was added after 48-72 hours upon confirmation of fibroblast outgrowth.

Primary Ai9 fibroblasts were cultured in DMEM + GlutaMAX (ThermoFisher) supplemented with 10% fetal bovine serum (Alstem) and 100U/mL of penicillin and streptomycin (Life Technologies). Cells were maintained at 37°C and 5% CO_2_ and passaged using standard cell culture techniques.

### Generation of neural progenitor cells (NPCs) from iPSCs and NPC culture

iPSCs (GM25256; NIGMS Human Genetic Cell Repository) were a gift from Bruce Conklin at the Gladstone Institute. Informed consent was provided for research use. All experimental procedures with this line were approved by Stanford’s Stem Cell Research Oversight (SCRO) panel.

iPSC-derived NPCs were generated by dual inhibition of SMAD signaling. First, embryonic bodies (EBs) were generated from iPSCs using the AggreWell 800 system (STEMCELL Technologies). On day 1, iPSC colonies were detached via Accutase (Innovative Cell Technologies) and collected by pipetting with N2/B27 Medium (DMEM/F12 (ThermoFisher) supplemented with 1x N-2 (ThermoFisher), 1x B27 without vitamin A (ThermoFisher), and 10µM Rock Inhibitor (Tocris)). 2e6 cells were plated per well of an AggreWell 800 plate in a total volume of 2mL. Cells were incubated at 37°C and 5% CO_2_ for 24 hours. On day 2, cells were fed with N2/B27 media supplemented with dual SMAD inhibitors (0.1μM LDN193189 (ThermoFisher) and 10μM SB431542 (ThermoFisher)) by removing 1mL of media and adding 1mL of N2/B27 media with 2x dual SMAD inhibitor. From day 3 to day 7, cells were fed every second day with N2/B27 media supplemented with 0.1μM LDN193189 and 10μM SB431542. On day 7, once iPSCs had aggregated to form EBs, they were plated onto Matrigel (Sigma) coated plates for another 7 days to form neural rosettes. During this time, cells were fed every second day with N2/B27 media supplemented with 0.1μM LDN193189 and 10μM SB43154. On day 14, neural rosettes were harvested using Neural Rosette Selection Reagent (STEMCELL Technologies) and plated in Matrigel-coated 6-well plates in NPC media (DMEM F12 (ThermoFisher) supplemented with 1x N-2 (ThermoFisher), 1X B27 without vitamin A (ThermoFisher), and 20ng/mL of FGF2 (ThermoFisher) in 0.1% Bovine Serum Albumin (Sigma)). For the next seven days, NPCs were fed every second day with NPC media.

NPCs were cultured in DMEM F12 (ThermoFisher) supplemented with 1x N-2 (ThermoFisher), 1x B27 without vitamin A (ThermoFisher), and 20ng/mL of FGF2 (ThermoFisher) in 0.1% Bovine Serum Albumin (Sigma). Cells were maintained at a density of ∼1-2e6 cells per well in a 6-well plate coated with Matrigel (Sigma), and media was changed every other day.

### Production of DARTs

Lenti-X cells were seeded in 15 cm dishes (Corning) at a density of 12.08e6 cells per dish. After approximately 16-18 hours, cells were transfected using TransIT-LT1 reagent (Mirus Bio) according to the manufacturer’s instructions. v2 and v3DARTs were produced by transfecting a plasmid mixture of mVSV-G (3.78µg), U6-sgRNA Gag-Cas9 (11.33µg), U6-sgRNA Gag-Pol (3.78µg), and scFv entry plasmid (11.33µg) in each 15 cm dish; for v1DARTs, 3.78µg of U6-sgRNA Gag-Cas9 and 11.33µg of U6-sgRNA Gag-Pol were used, instead. For other production scales, plasmid and producer cell amounts were scaled by plate surface area. Unless otherwise stated, DARTs were produced in 15 cm dishes. A 1x dose of DARTs is defined as the amount of VLPs generated in 2mL of producer cell media.

### Production of DART LV

Lenti-X cells were seeded in 6-well plates (Corning) at a density of 1.2e6 cells per dish (Corning). After approximately 16-18 hours, cells were transfected using TransIT-LT1 reagent (Mirus Bio) according to the manufacturer’s instructions. DART LV was produced by transfecting a plasmid mixture of psPAX2 (1.3µg), pMD2.G with mVSV-G and anti-FITC scFv or mSA (pSLQ11876 or pSLQ11875, respectively; 555ng), and an mCherry-containing transfer plasmid (1.8µg) in each well.

### Production of MMLV base and gene editor DARTs

Lenti-X cells were seeded in 15 cm dishes (Corning) at a density of 9.67e6 cells per dish. After approximately 16-18 hours, cells were transfected using TransIT-LT1 reagent (Mirus Bio) according to the manufacturer’s instructions. Base and gene editor DARTs were produced by transfecting a plasmid mixture of mVSV-G (0.8ug), Gag-Pol (6.5ug), Gag-ABE or Gag-Cas9 (2.2ug), gRNA (8.5ug), and scFv entry plasmid (2.3 or 0.8ug). For other production scales, plasmid and producer cell amounts were scaled by plate surface area.

### DART isolation and transduction for *in vitro* studies

48 hours post-transfection, DARTs were harvested and filtered through 0.45µm syringe filters (Fisher Scientific), and 5x Lenti-X solution (Alstem) was added for overnight incubation at 4°C. The following day, DARTs were concentrated via centrifugation: we performed a 30-minute spin at 1,500xg and 4°C, aspirated the supernatant, ran another 5-minute spin at 1,500xg and 4°C, and aspirated the supernatant again. DARTs were then resuspended in the appropriate cell culture media in an appropriate volume, never exceeding 100x concentration of the harvested volume. For all experiments, a 1x dose was defined as 2mL of harvested supernatant. If not used for transduction the day after harvesting, DARTs were flash frozen in ethanol and dry ice and stored at −80°C after concentration and resuspension. DART LV was harvested in the same manner, except harvesting occurred 24 hours post-transfection, instead of 48 hours post-transfection.

We employed two different methods for transduction with DARTs: 1) “pre-staining” cells with antibody prior to transduction or 2) conjugating antibodies to DARTs prior to transduction. For the former, cells were spun down and resuspended in 99uL of DPBS and 1uL of the targeting antibody. Pre-stained cells were incubated in the dark with rocking for 20-30 minutes. Afterwards, cells were washed twice via centrifugation and transduced with DARTs. For the latter, FITC-conjugated or biotinylated antibodies diluted to 10x the desired concentration were added to DARTs at a 1:10 ratio and incubated at room temperature in the dark for 30 minutes. The virus-antibody mixture was subsequently added to cells.

### Endocytosis inhibitor experiments

HEK293T cells and CD19+, GFP+ HEK293T cells were seeded in 48-well plates (Corning) at 3e4 cells per well. Approximately 24 hours later, prior to transduction with v3DARTs, cells were incubated for 60 minutes with dynasore (MedChem) or EIPA (MedChem) (which were diluted in DMEM to the desired concentrations prior to use). Heparin (Sigma) was added to v3DARTs post-isolation at the specified concentrations, with incubation for 15-30 minutes prior to αCD19 antibody conjugation. Cells were then transduced with v3DARTs as described above. Media was changed the following morning.

### DART isolation via EXODUS for characterization and *in vivo* studies

48 hours post-transfection, DARTs were harvested and filtered through 0.45µm syringe filters (Fisher Scientific), then shipped to collaborators at the Wang Lab (UC Davis) overnight on ice. DARTs were isolated from supernatants using the EXODUS H-600 (Exodus Bio, USA). 30nm pore size nanoporous anodic aluminum oxide filters with a large-size exosome isolation device (EID) designed for processing volumes of 50-250mL were used. DART isolation was performed following EXODUS H-600 manufacturer’s instructions. Harvested media was concentrated down to 250mL using 100 kDa Amicon Ultra filters (Millipore Sigma, USA). The sample was filtered through a 0.22µm PVDF syringe filter (Millipore Sigma, USA) to remove large particles and run on the EXODUS H-600 using the *CellCulture.Strong-MA.A03* default program. After isolation, DARTs were recovered from the EID and aliquoted into 1.5mL Eppendorf tubes. All aliquots were stored at −80°C. For *in vivo* studies, DARTs were conjugated to antibodies prior to transduction as described above.

### Immunostaining and flow cytometry for *in vitro* studies of cultured cells

For cell lines and primary human cells, immunostaining was used to quantify knockout of the targeted gene (e.g., TRAC, B2M). For primary T cells, immunostaining was also used to determine CD4 and CD8 positivity. mCherry fluorescence was used to approximate editing of the tdTomato locus in Ai9 fibroblasts. EGFP fluorescence was used to approximate editing in the base editor reporter cell line.

Cells were washed with DPBS or FACS buffer (DPBS supplemented with 2% FBS and 1 mM EDTA (Gibco)) prior to flow cytometry. Cells were stained in the dark for 20-30 minutes with near-IR (780) live-dead stain (ThermoFisher). After staining, cells were washed once with DPBS or FACS buffer and subsequently stained in the dark for 20-45 minutes with fluorophore-conjugated antibodies. After staining, cells were washed twice with DPBS or FACS buffer and resuspended in a final concentration of 150uL FACS buffer. Flow cytometry was performed on a CytoFLEX S (Beckman Coulter). For all samples, at least 1e4 cells were gated within the population of interest (Live/Single Cells). FlowJo (FlowJo LLC) software was used for data analysis.

### Amplicon sequencing to assess genome editing

Sanger sequencing was used to detect on-target genome editing in NPCs. Genomic DNA was extracted using QuickExtract DNA Extraction Solution (BioSearch) according to the manufacturer’s instructions. CloneAmp DNA polymerase (Takara Bio) was used to amplify the Cas9 sgRNA target site. PCRs were purified using NucleoSpin Gel and PCR Clean-up Columns (Machereney-Nagel) and sent to GENEWIZ from Azenta Life Sciences for Sanger sequencing. Editing efficiency was computed using publicly-available ICE software (https://ice.editco.bio/<u>#/</u>).

### Cas9 protein quantification

Cas9 concentration in v1, v2, v3DARTs, and aCD3 VLPs was determined using the SpCas9 ELISA kit (Cell Biolabs), according to the manufacturer’s instructions. DARTs were produced in 6-well plates and concentrated as described above. A 1/4x dose of each was lysed in 750uL lysis buffer (RIPA buffer (ThermoFisher) with 1x Protease Inhibitor Cocktail (Sigma) and 1mM DTT) and centrifuged for 10 minutes at 10,000xg and 4°C. 100uL of lysed DARTs were added per well of the anti-Cas9 antibody coated plate. Following substrate development, the optical density (OD) of the plate was measured at 450nm using a Synergy H1MG fluorescent plate reader (BioTek). The concentration of Cas9 was calculated using the supplied Cas9 standard curve. To calculate amount of Cas9 per DART particle, particle counts were estimated using NTA data for the corresponding formulations.

### Western blot analysis

DARTs were produced in 6-well plates (Corning) and concentrated as described above. Samples were resuspended in 50uL (40x concentration) in Dulbecco’s Phosphate Buffered Saline (DPBS) (Gibco). Samples were then mixed with 200uL lysis buffer (RIPA buffer (ThermoFisher) with 1x Protease Inhibitor Cocktail (Sigma) and 1mM DTT) and centrifuged for 10 minutes at 20,000xg to remove any leftover debris. Supernatant was transferred to a new tube, and 4x LDS Sample Buffer (Invitrogen) was added to each sample prior to boiling for 10 minutes at 95°C. 4.5uL/sample was loaded into a 4-12% Bis-Tris Gel (ThermoFisher). Gels were run in MOPS buffer (Invitrogen) at 200V for 40-50 minutes. The proteins were transferred to a PVDF membrane using an iBlot2 Gel Transfer Device (ThermoFisher) set at 20V for 5 minutes and 20 seconds. Membranes were then cut at approximately 65kDa to separate the high molecular weight and low molecular weight proteins.

The two halves were blocked for 30 minutes at room temperature with rocking in a blocking buffer (3-5% non-fat milk in 1x tris-buffered saline with 0.1% Tween-20 (TBST). After blocking, membranes were washed in 1x TBST three times for 5 minutes. HRP-conjugated mouse anti-FLAG antibody (Sigma) and purified rabbit anti-HIV p24 antibody (Sigma) were diluted in 1% non-fat milk in 1x TBST 1:10,000 and 1:2,500, respectively. The anti-FLAG antibody was added to the high molecular weight portion of the membranes, and the anti-HIV p24 antibody was added to the low molecular weight portion. They were then incubated overnight at 4°C.

The following morning, blots were washed three times for 15 minutes with rocking at room temperature in 1x TBST. Goat anti-rabbit HRP diluted 1:5,000 in 1% non-fat milk in 1x TBST was added to the p24-stained membrane and incubated for 1 hour at room temperature with rocking. After secondary staining, p24-stained membranes were again washed three times for 15 minutes with rocking at room temperature in 1x TBST. The HRP was developed using Azure Biosystems Radiance (Azure Biosystems) according to the manufacturer’s instructions and imaged with Azure 300 Chemiluminescence Gel Imager (Azure Biosystems).

### Characterization of DARTs by nanoparticle tracking analysis

Nanoparticle Tracking Analysis (NTA) was performed to quantify DARTs’ concentration and size distribution. The ZetaView x30 Next Generation Nanoparticle Analyzer (Particle Metrix) was used to characterize all formulations. DARTs were diluted 200,000x into 4mL of Ultrapure Water (Gibco) and injected into the NTA. Size and zeta potential measurements were collected at 22°C.

### Cryo-electron microscopy

Cryogenic transmission electron microscopy (cryo-EM) was used for direct visualization of DARTs. To prepare samples for cryo-EM, copper grids (200 mesh, 1.2µm holes; Ted Pella Inc) were glow discharged for 30 seconds at 25mA using a Pelco EasiGlow system (Ted Pella Inc). A 4uL aliquot of DARTs was applied to the carbon side of the grid, blotted for 5 seconds in the humidity chamber, incubated for 30 seconds, and plunge-frozen into precooled liquid ethane in an EM GP2 Automatic Plunge Freezer (Leica Microsystems). This process results in the vitrification of DARTs in a thin layer of amorphous ice to preserve them in their native state and protect against radiation damage during imaging.

The samples were then imaged using a Glacios 2 electron microscope (ThermoFisher) equipped with an autoloader with an accelerating voltage of 200kV for high resolution grid screening and data collection. Low dose mode was used during image acquisition to minimize radiation damage of the DARTs samples. Images were obtained with a GATAN K3 direct electron detector at 11,000x and 45,000x magnification with a defocus value between −2µm and −5µm. The total accumulated dose per image did not exceed 40e/Angstrom^2.

### Super resolution microscopy

Single molecule localization microscopy was used to obtain super resolution images of DARTs for analysis of surface marker expression and quantification of Cas9 encapsulation efficiency. Direct stochastic optical reconstruction microscopy (dSTORM) of individual DARTs was performed using a temperature-controlled Nanoimager S Mark II microscope (Oxford Nanoimaging) equipped with a 100x, 1.4NA oil immersion objective. The EV Profiler Kit (version 1; Oxford Nanoimaging), was used per manufacturer’s instructions. Surface marker analysis of DARTs samples was performed using biotin (5-fluorescein) conjugate (Sigma) or rat anti-mouse FITC-CD184 (BD Pharmingen), FLAG tag (DYKDDDDK)-AlexaFluor 594 antibody (ThermoFisher), and lipophilic Memglow700 dye (Cytoskeleton Inc).

Image acquisition on the Nanoimager was performed in total internal reflection fluorescence (TIRF) mode with a 54° illumination angle. Laser powers were calibrated for each use of the microscope along with channel mapping calibration to align channel data with a maximum 10nm standard deviation for calculated point spread functions. 1,000 frames of data were acquired with a 30ms exposure per channel (3,000 frames total) using NimOS software (v1.19.7; Oxford Nanoimaging). Raw localization images were uploaded to CODI for cloud storage and automated image analysis. Briefly, localization images were drift corrected, filtered by photon count and precision, clustered, and counted for population statistics and relative fluorophore abundance analysis.

For analysis of dSTORM data, clusters positive in only a single channel were excluded from analysis. Single-positive clusters were excluded to minimize contributions from nonspecific fluorescence, free fluorophore/antibody aggregates, free dye, membrane or particle fragments, and other non-particle-associated structures. Clusters positive in two or three channels remained in the analysis pool and are termed “identified particles” in the main text; these were used for the analysis performed in **Fig. 4d,e**. For a full report on single-, double-, and triple-positive clusters, see **SI Fig. 2**.

### Animal studies

All animal procedures were approved by The University of California, Davis (UCD) institutional animal care and use committee (IACUC). All facilities used during the study period were accredited by the Association for the Assessment and Accreditation of Laboratory Animal Care International (AAALAC). Ai9 transgenic mice were purchased from The Jackson Laboratory (Jackson stock No. 007909). Ai9 mice were time-mated and bred on-site at UCD. The *in utero* injections were performed as previously described^82–84^. Pregnant mice at embryonic day 15.5 (E15.5) were anesthetized with 5% isoflurane, followed by 2% isoflurane for maintenance, and placed in the supine position on a heating pad. After a 2cm midline abdominal incision was made, the uterus was gently exteriorized using cotton tipped applicators. The number of fetuses on either side of the uterine bifurcation was noted, along with the viability status of each fetus. A glass pipette, which had been pulled to an opening diameter of 50-70μm, was loaded with 2μL of Exodus-concentrated DARTs alone or conjugated to antibody at the specified concentration as described above. PBS was injected as a control. The glass pipette was inserted through the fetal skull and brain tissue into one of the lateral cerebral ventricular spaces. The uterus was then gently interiorized back into the abdomen using cotton tipped applicators and the abdomen was filled with approximately 1mL of 37°C saline to avoid dehydration. The maternal laparotomy was then sutured closed in 2 layers using absorbable 4-0 vicryl. The mouse was given a subcutaneous injection of 0.05 mg/kg of buprenorphine for pain management every 12 hours for the next 48 hours. After the *in utero* injections, the pregnant dams continued to gestate until the offspring were either delivered via cesarean section at E19.5 or until natural and full-term delivery occurred.

### Isolation of Embryonic Neural Stem Cells from Mouse Fetal Brain

Embryonic neural stem cells (NSCs) were isolated from the fetal brains of E15.5 Ai9 mouse embryos. Pregnant female mice were euthanized following institutional animal care and use guidelines, and the uterine horns were harvested in sterile ice-cold Hank’s Balanced Salt Solution (HBSS, ThermoFisher). Embryos were dissected from the amniotic sac, and the heads were removed under a dissection microscope. The cerebral cortices were carefully isolated from the meninges and transferred into a sterile HBSS containing 10% penicillin-streptomycin (ThermoFisher). The tissue was gently minced using fine scissors and enzymatically dissociated in 0.05% Trypsin-EDTA (ThermoFisher) at 37°C for 10 minutes with occasional gentle mixing. The enzymatic reaction was stopped by adding an equal volume of trypsin inhibitor, followed by centrifugation at 300xg for 5 minutes. The cell pellet was resuspended in NSC culture medium consisting of Neurobasal Medium (ThermoFisher) supplemented with 2% B27 without Vitamin A (Gibco), 20ng/mL epidermal growth factor (EGF, Advent Bio), 20ng/mL basic fibroblast growth factor (bFGF, Advent Bio), 1 U/ml heparin (STEMCELL Technologies), 1% L-Glutamine (Gibco), 1% MEM non-essential amino acids (ThermoFisher), 0.1mM b-mercapoethanol (Sigma) and 1% penicillin-streptomycin (ThermoFisher). The dissociated cells were filtered through a 40µm cell strainer (Falcon) to remove debris and then plated in culture dishes (Corning) at a density of 1e5 cells/mL. Cells were maintained in a humidified incubator at 37°C with 5% CO_2_, and neurosphere formation was monitored daily. The medium was refreshed every two to three days. After 5-7 days in culture, neurospheres were collected, dissociated using 0.05% Trypsin-EDTA, and replated for expansion or downstream assays.

### Flow cytometry for *in vivo* studies

The transduction rate of the fetal brain was measured by flow cytometry based on the percentage of cells positive for tdTomato following *in utero* treatment. Briefly, freshly dissected brains were minced and then digested with 600μL of 1 mg/mL collagenase type I (Gibco) at 37°C, 5% CO_2_ for 20 minutes in 1.5mL Eppendorf tubes. Following this, the collected cell suspension was neutralized with 10% fetal bovine serum (FBS) and kept on ice. Additional 1% collagenase 1 solution was added to the remaining tissue and incubated in the same conditions for an additional 20 minutes. A single-cell suspension was obtained by filtering the resulting cell suspension solutions through a 70μm Nylon cell strainer (Falcon). The Attune NxT Flow Cytometer (ThermoFisher) was used for performing flow cytometry, and FlowJo software (FlowJo LLC) was used for data analyses.

### Histology and immunofluorescence staining

Adult mice were perfused with PBS and 4% paraformaldehyde (PFA). Dissected fetal brains were fixed in 4% PFA for 24 hours, dehydrated with 15% sucrose for 24 hours, followed by 30% sucrose for 24 hours, before being embedded in O.C.T. (Fisher Healthcare; Scigen Scientific Gardena; 4585). Serial 10μm thickness sections were made using a Cryostat (Leica CM3050S) before sections were collected onto Matsunami Glass microscope slides. For immunofluorescence staining, tissue sections were washed gently and thoroughly with PBS before undergoing heat antigen retrieval with 10mM sodium citrate buffer pH 6.0. Sections were then blocked with 5% Bovine Serum Albumin (BSA) in PBS at room temperature for 1 hour. Sections were stained with primary antibody solutions at 4°C overnight. Primary antibodies were used at the following dilutions: 1:250 for tdTomato (MyBioSource, MBS448092), 1:100 for NeuN (Cell Signaling Technologies, D4G40, 24307S), 1:250 for Sox2 (Cell Signaling Technologies, L1D6A2; 4900S), and 1:400 for Nestin (Cell Signaling Technologies, 42791). Sections were stained with the corresponding secondary antibody solution at room temperature for 30 minutes. The dilution of secondary antibodies was 1:250. Sections were then stained with a 1:5,000 dilution of DAPI at room temperature for 5 minutes. Finally, slides were mounted with Prolong Diamond Antifade Mountant (Invitrogen) and imaged by ImageXpress Pico automated cell imaging system (Avantor) or Zeiss Observer Z1.

### FIJI-based image quantification

To quantify target-cell specificity and percentage of edited Sox2^+^ cells, FIJI was used to analyze confocal images of brain sections from nanoparticle-treated fetuses. Regions containing discernable editing were used for analysis; this included 2 regions from one brain section for CD56-treated DARTs (**Fig. 6e**, **SI Fig. 5f**) and 1 region for DARTs alone (**Fig. 6g**). First, the raw images were split into separate channels and individually thresholded to maximize the distinction between adjacent nuclei. The image calculator AND function was used to create masks where DAPI and Sox2, DAPI and tdTomato, or all three signals were present. For DAPI^+^/Sox2^+^ cells, the count was performed using the Analyze Particles function, with a size distinction of 3-infinity and any degree of circularity. Counts for the other two groups were performed by hand.

“Target-cell specificity” figures were computed as the number of DAPI^+^/Sox2^+^/tdTomato^+^ cells divided by the number of DAPI^+^/tdTomato^+^ cells, reported as a percent value. The percentage of edited Sox2^+^ cells were computed as the number of DAPI^+^/Sox2^+^/tdTomato^+^ cells divided by the number of DAPI^+^/Sox2^+^ cells.

### Statistical analysis

Statistical tests for significance between groups were performed in GraphPad Prism 10. Details on all statistical tests can be found in the figure legends. When applicable, all P-values were corrected for multiple comparisons testing. In figures, all statistically-significant P-values were reported, with asterisks defined as follows: ∗*P* <0.05, ∗∗*P* <0.01, ∗∗∗*P* <0.001, ∗∗∗∗*P* <0.0001. Data points were excluded only when a clear technical failure or predefined quality-control criterion indicated that the measurement was unreliable. Potential outliers were identified and verified using the ROUT method in GraphPad Prism 10 (Q = 5%).

## Acknowledgements

The authors thank all members of the Lei (Stanley) Qi lab for facilitating experiments and helpful discussions. We thank C. Chen and T. Wachsmann from the Qi Lab for assistance with T cell isolation and culture; Y. Ma and X. Chen from the Qi Lab for productive subgroup discussions; M. Fu and E. Li from the Qi Lab for generation of induced pluripotent stem cell-derived neural progenitor cells; M. Fu from the Qi Lab for facilitation of *in vivo* experiments; P. Desai from the Wang Lab for expert assistance with cryo-electron microscopy; S. Wang from Stanford Ophthalmology for generous guidance in *in vivo* experimentation; Y. Cheng for cloning assistance during her graduate rotation. G.R. acknowledges support from the NIH Stanford Graduate Training Program in Biotechnology (T32GM141819). I.R.G. acknowledges support from the Stanford Graduate Fellowship and the Sarafan ChEM-H Chemistry/Biology Interface Training Program. N.A. acknowledges support from the CIRM Stem Cell and Gene Therapy Training Program 2.0. D.W. acknowledges support by the NIH Cardiovascular Biology T32 Predoctoral Fellowship (HL086350). A.W. acknowledges support by the National Institutes of Health, the California Institute for Regenerative Medicine (CIRM), and Shriners Children’s. L.S.Q. acknowledges support from the National Science Foundation, the National Institutes of Health, the Chau Li Research Fund, the Weill Cancer Hub West, and the Research to Prevent Blindness (RPB). This work was supported by Shriners Children’s (85400-NCA-24), the California Institute for Regenerative Medicine (DISC2-14097), the National Institute on Aging (R21AG077193), National Cancer Institute (R01CA266470), National Institute of Neurological Disorders and Stroke (R01NS144390), National Human Genome Research Institute (R21HG013133), NIH Director’s Pioneer Award (DP1NS137219), the Weill Cancer Hub West through Project PROMISE, and the Research to Prevent Blindness (RPB) Stein Innovation Award. L.S.Q. is a Chan Zuckerberg Biohub Investigator. Its contents are solely the responsibility of the authors and do not necessarily represent the official views of the National Institutes of Health.

## Contributions

M.C., P.F., and L.S.Q. conceived of the foundational idea for the work. G.R., I.G., and L.S.Q. discussed the primary research approach and strategy. G.R. and I.G. performed *in vitro* experiments in Figures 1, 2, 3, and 5 and produced all VLP batches for *in vitro* and *in vivo* experimentation. M.C., P.F., and G.R. performed *in vitro* experiments in Figure 6. G.R., I.G., N.A., K.G., D.W., L.S.Q., and A.W. discussed the *in vivo* research approach and strategy. N.A. and K.G. performed all *in vivo* work and whole-brain flow cytometry. N.A., K.G., D.W., and S.X. performed nanoparticle tracking assays. D.W. performed direct stochastic optical reconstruction microscopy and cryogenic transmission electron microscopy experiments and analyzed the data. G.R. prepared the manuscript and figures with the assistance of I.G. L.S.Q. made significant revisions to the manuscript. G.R., I.G., M.C., and P.F. were supervised by L.S.Q., and N.A., K.G., D.W., and S.X. were supervised by A.W. All authors have read and approved the manuscript.

## Ethics Declarations

M.C., P.B.F., and L.S.Q are co-inventors of a patent filed by Stanford University on related work (63/405,720). L.S.Q. is a scientific founder of Epicrispr Biotechnologies. M.C. and P.B.F. are founders of Enoda Cellworks.

## SI Figures

**SIFigure 1:**
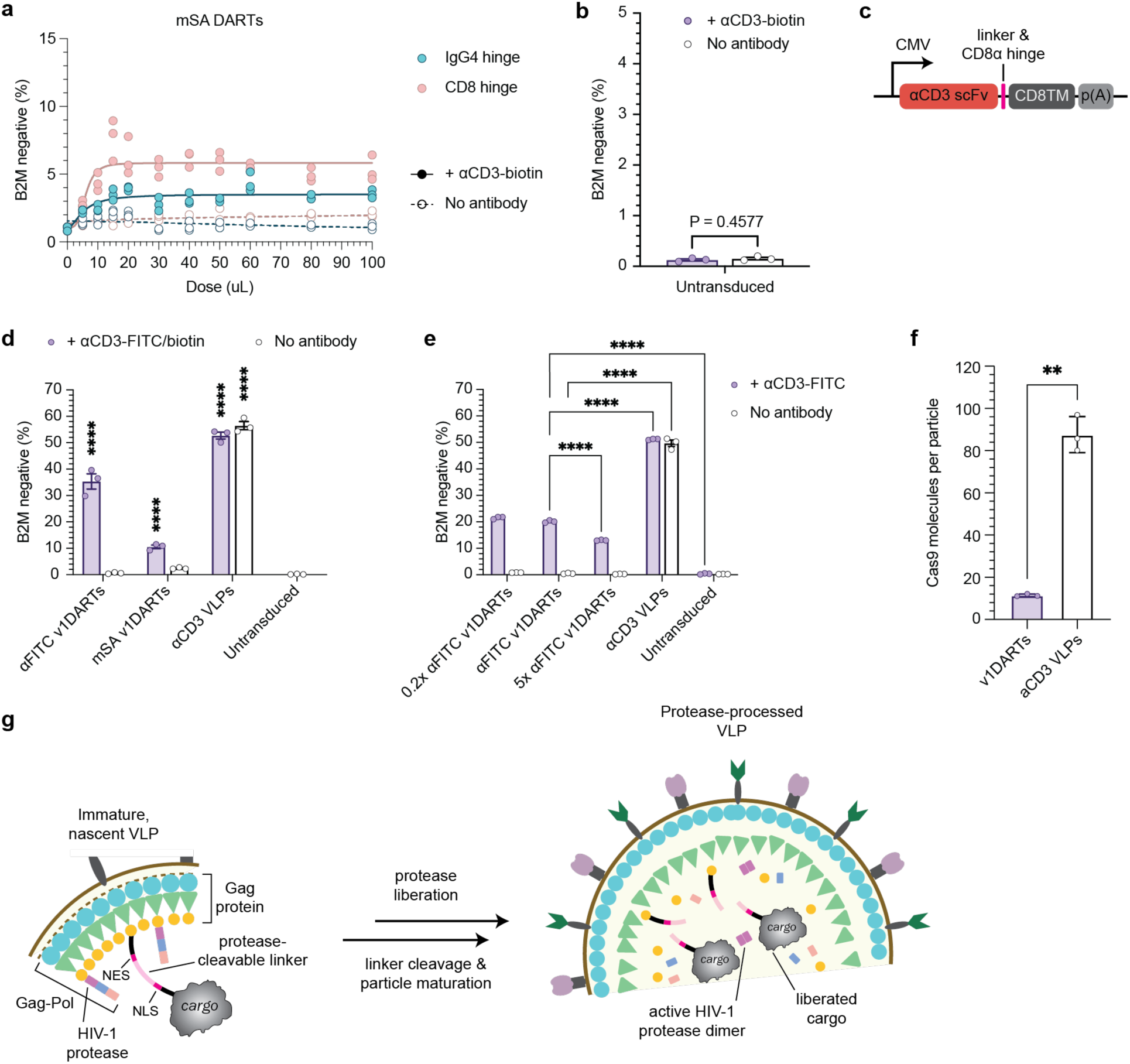
Supporting information for optimization of DARTs and comparison with membrane-integrated binders. **a,** Architecture optimization of mSA DARTs, as in Fig. 1c. DARTs were concentrated 5x prior to transduction. All replicates are plotted and curves were fitted via sigmoidal four-parameter logistic regression. **b,** Control experiment to determine if the pre-staining protocol affected B2M expression without the addition of DARTs. Cells were stained with αCD3-biotin antibody, excess was removed, then cells were resuspended in media; B2M knockout was measured by flow cytometry 6 days later. **c,** Schematic of the construct used to produce VLPs with an αCD3 scFv decorating the outer membrane (termed αCD3 VLPs). **d,** Comparison of αFITC and mSA v1DARTs to αCD3 VLPs, all packaging sgB2M. Jurkat cells were transduced with a 1x dose of all three formulations using the pre-staining method. B2M knockout was quantified by flow cytometry 6 days post-transduction. Statistical comparison is to the untransduced condition. **e,** αFITC v1DARTs were produced with varied amounts of the αFITC scFv plasmid with the CD8α hinge. Transduction conditions were as in **d**. All conditions were compared to one another within +/− antibody groups. **f,** Quantification of Cas9 molecules per particle in αFITC v1DARTs vs αCD3 VLPs. Cas9 was quantified by ELISA and normalized to particle concentration obtained via nanoparticle tracking analysis (NTA). Statistical comparison was performed via Welch’s t test. For all data, n=3 technical replicates per condition. Bar graphs are mean ± s.d. Unless otherwise stated, statistical analysis was conducted via two-way ANOVA. For all plots: *, P<0.05; **, P<0.01; ***, P<0.001; ****, P<0.0001. **g,** Schematic of the cargo liberation and maturation process in DARTs. During the particle’s maturation process, the protease dimerizes, activates, and cleaves at the recognition sequence, liberating the cargo from Gag and leaving it free to enter the nucleus after VLP infection. Antibody clones used for transduction: αCD3, HIT3a. Antibody clone used for staining: αB2M, 2M2. Abbreviations: B2M, β-2 microglobulin; scFv, single-chain variable fragment; CD8TM, CD8α transmembrane domain; p(A), poly-A tail; NES, nuclear export sequence; NLS, nuclear localization sequence.

**SI Figure 2:**
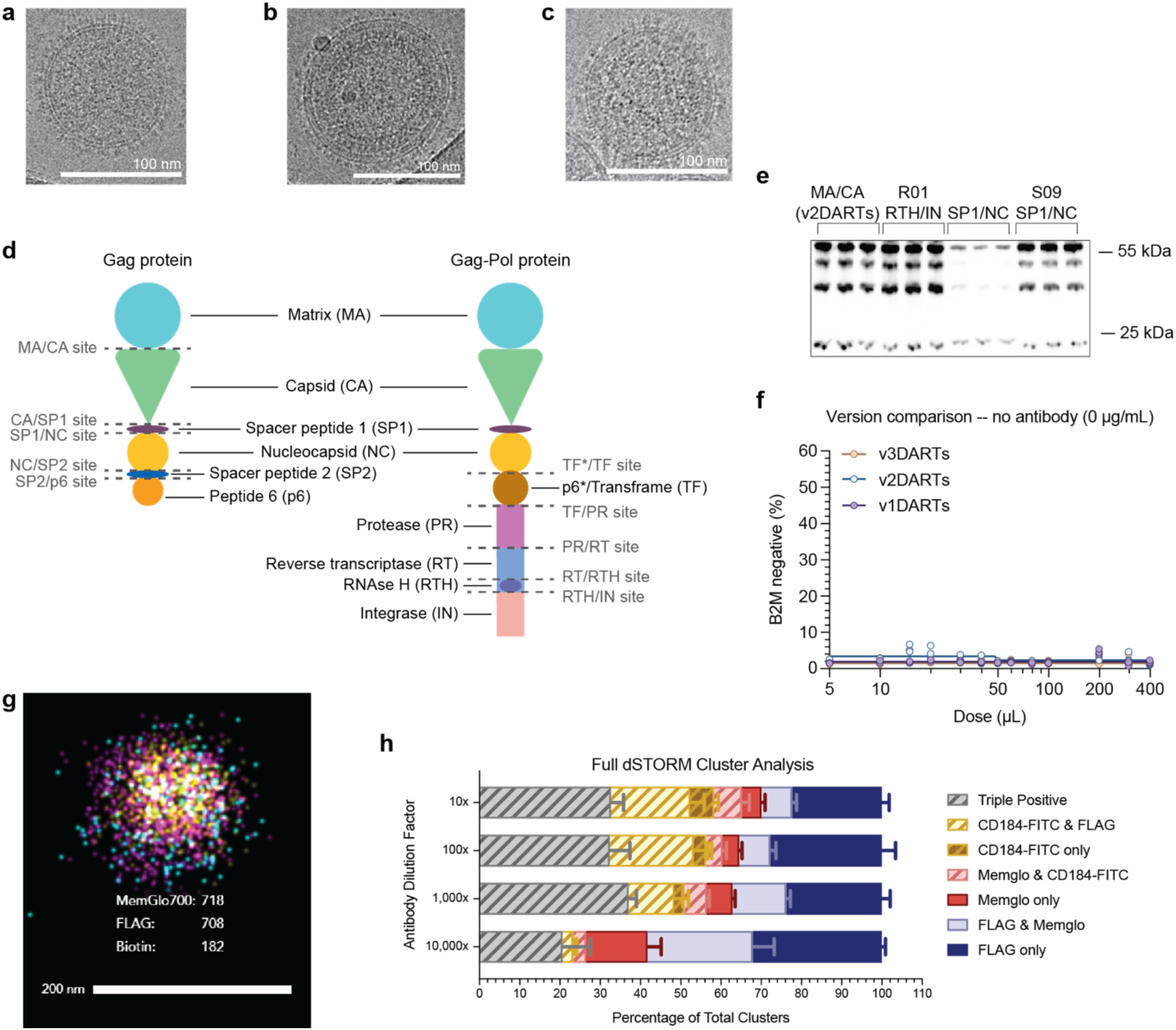
Additional information regarding DARTs’ structure, antibody-independent activity, antibody conjugation efficiency, and Cas9 encapsulation fidelity. **a**-**c,** Additional cryo-EM images of αFITC v2DARTs. Images depict particles with undefined (**a**), immature (**b**), and mature (**c**) inner membranes. Scale bars = 100 nm. **d,** Schematic of the individual protein components (labels in black) that make up the Gag and Gag-Pol polyproteins, and the ten protease recognition sites (labels in gray) that separate them. **e,** Additional western blot for p24, as in **Fig. 3f**. Data replicates and supports the earlier findings. **f,** As in **Fig. 3j-l**, αFITC v1, v2, and v3DARTs are compared across doses in the absence of targeting antibody. All replicates are plotted and curves were fitted via sigmoidal four-parameter logistic regression. **g,** Additional, representative image of v2DARTs captured via dSTORM. **h,** Related to **Fig. 4d,e**. αCD184-FITC was serially diluted and used to stain αFITC DARTs prior to dSTORM imaging. Memglow was used to stain the membranes of DARTs, while FLAG stained for the presence of Cas9. The raw cluster information is plotted as a parts of whole graph, but single-positive clusters are not eliminated from the analysis. Data was collected as one lane per sample and averaged across four fields of view per lane. Data are all mean ± s.e.m. Antibody clone used for transduction: αCD3, HIT3a. Antibody clone used for staining: αB2M, 2M2. Antibody clone used for imaging: αCD184, 2B11.

**SI Figure 3:**
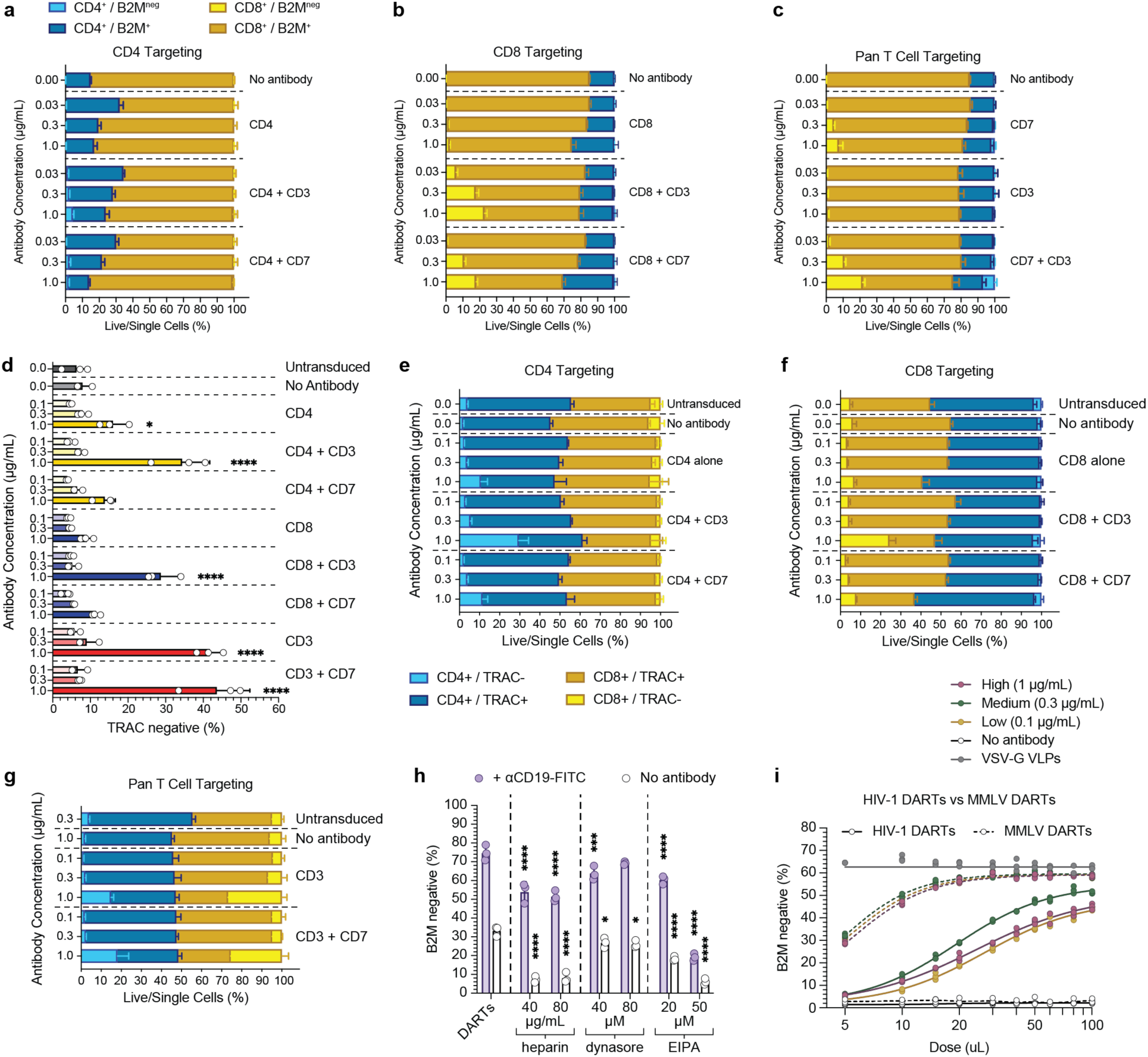
Additional characterization of DARTs’ specificity, off-target activity, and editing efficiency across VLP scaffolds. **a**-**c,** A 4x dose of αFITC v3DARTs was targeted towards one primary human T cell donor via a panel of antibodies. The raw data used to produce the graphs shown in **Fig. 4b,d**, and **f** are shown as part of whole plots in **a,b**, and **c**, respectively. **d,** A 4x dose of αFITC v2DARTs carrying sgTRAC was targeted towards a different primary human T cell donor via a panel of antibodies. The depicted values are representative of the entire pool of T cells, irrespective of T cell subtype. This data is a repeat of that shown in Fig. 4a, but a different gene was targeted and a different blood donor was used. Each group was compared with the no-antibody condition using two-way ANOVA. **e**-**g,** Parts of whole CD4^+^/CD8^+^ T cell population breakdown for the data shown in **d**. Plots depict the cell-type specificity of v2DARTs directed towards CD4^+^ (**e**) and CD8^+^ (**f**) T cell subtypes, and the equitable transduction that occurs when neither group is directly targeted (**g**). **h,** Raw B2M knockout data used to produce Fig. 5k. The inhibitor-dependent change in B2M knockout, plotted in Fig. 5k, was calculated using the DARTs only group (left). The plotted statistical significance compares each condition to the DARTs only condition with the same antibody treatment via two-way ANOVA. The two-way ANOVA revealed significant effects of inhibitor treatment (P < 0.0001) and antibody presence (P < 0.0001), as well as a significant interaction between these factors (F(6,28) ≈ 25, P < 0.0001), indicating that the effects of the inhibitors depended on whether antibody was present. **i,** Overlay of curves from **Fig. 3i** and Fig. 6b, comparing HIV-1-derived αFITC v3DARTs to MMLV DARTs. For all experiments, n=3 technical replicates per condition. Readout was performed via flow cytometry 6 days post-transduction. Data are all mean ± s.d. For all plots: *, P<0.05; **, P<0.01; ***, P<0.001; ****, P<0.0001. Antibody clones used for transduction: αCD4, SK3; αCD3, HIT3a; αCD7, CD7-6B7; αCD8, RPA-T8; αCD19, 4G7. Antibody clones used for staining: αB2M, 2M2; TRAC, IP26; αCD4, SK3; αCD8, RPA-T8. Abbreviations: B2M, β-2 microglobulin; TRAC, T cell receptor α constant.

**SI Figure 4:**
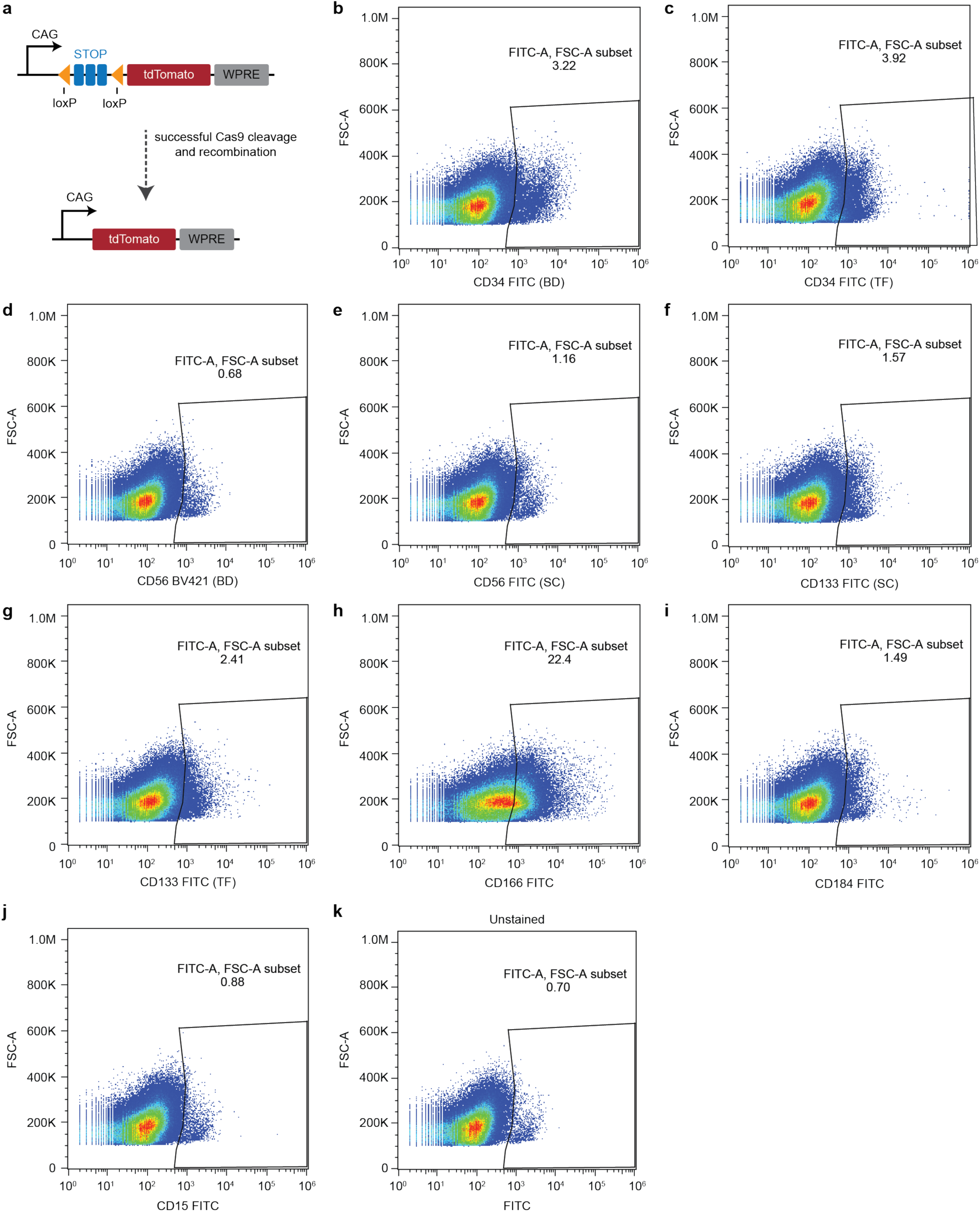
E15.5 fetal mouse brain antigen characterization. **a,** Basic schematic of how Cas9 editing with an sgRNA targeting LoxP (sgLoxP) can remove a stop cassette and induce tdTomato signal in the Ai9 mouse model. In Ai9 mice, this cassette is integrated into the *ROSA26* safe harbor locus. **b**-**j,** E15.5 Ai9 fetal mouse brains were harvested, digested, and stained with a panel of ten fluorophore-conjugated antibodies. Flow cytometry plots are shown from the nine that were not selected for *in vivo* experimentation. Parentheses following the x-axis fluorophore-conjugated antibody label indicate the antibody vendor (BD, BD Biosciences; TF, ThermoFisher; SC, Santa Cruz Biotechnology). **k,** Unstained control sample for comparison to **b**-**j**. Abbreviations: WPRE, Woodchuck Hepatitis Virus Posttranscriptional Regulatory Element; BD, BD Pharmingen; TF, ThermoFisher; SC, Santa Cruz Biotechnology.

**SI Figure 5:**
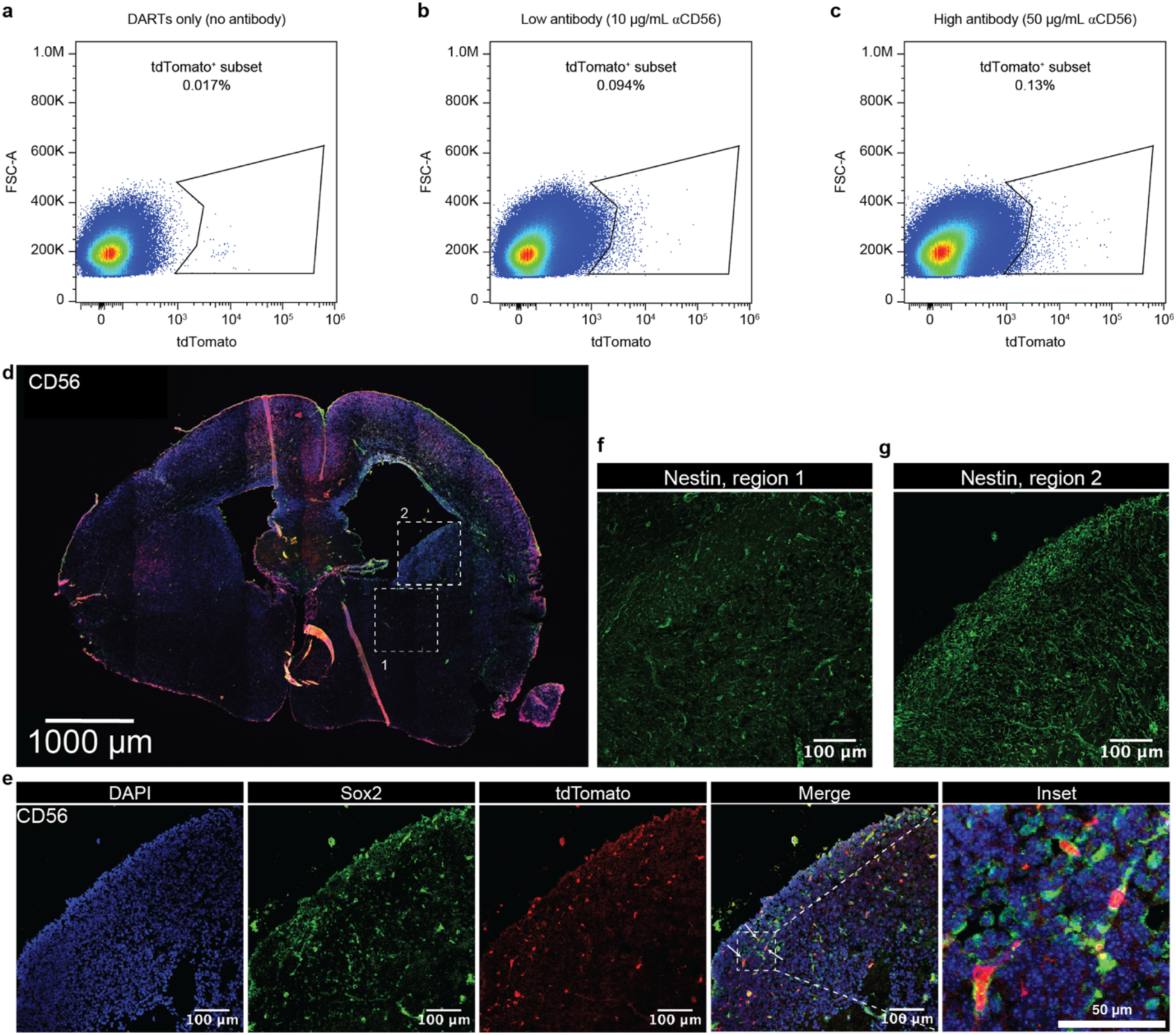
Whole-brain flow cytometry data and additional tissue section insets. **a-c,** Additional set of flow plots from representative no antibody (**a**), low antibody (**b**), and high antibody (**c**) samples plotted in Fig. 7c. **d,** Preserved brain section from an Ai9 mouse fetus treated with DARTs targeted by αCD56-FITC at 50 µg/mL. The image is as in Fig. 7g, with a second region highlighted for high-resolution analysis. Scale bar = 1 mm. **e,** Higher magnification of boxed region 2 from **d**. Arrows indicate highlighted tdTomato^+^/Sox2^+^ cells depicted within the inset. **f,g** Higher magnification of nestin staining from boxed regions 1 (**f**) and 2 (**g**) from **d**. (**e-g**) Scale bars = 100 µm, inset scale bar = 50 µm. Antibody clone used for transduction: αCD56, polyclonal.

**SI Figure 6:**
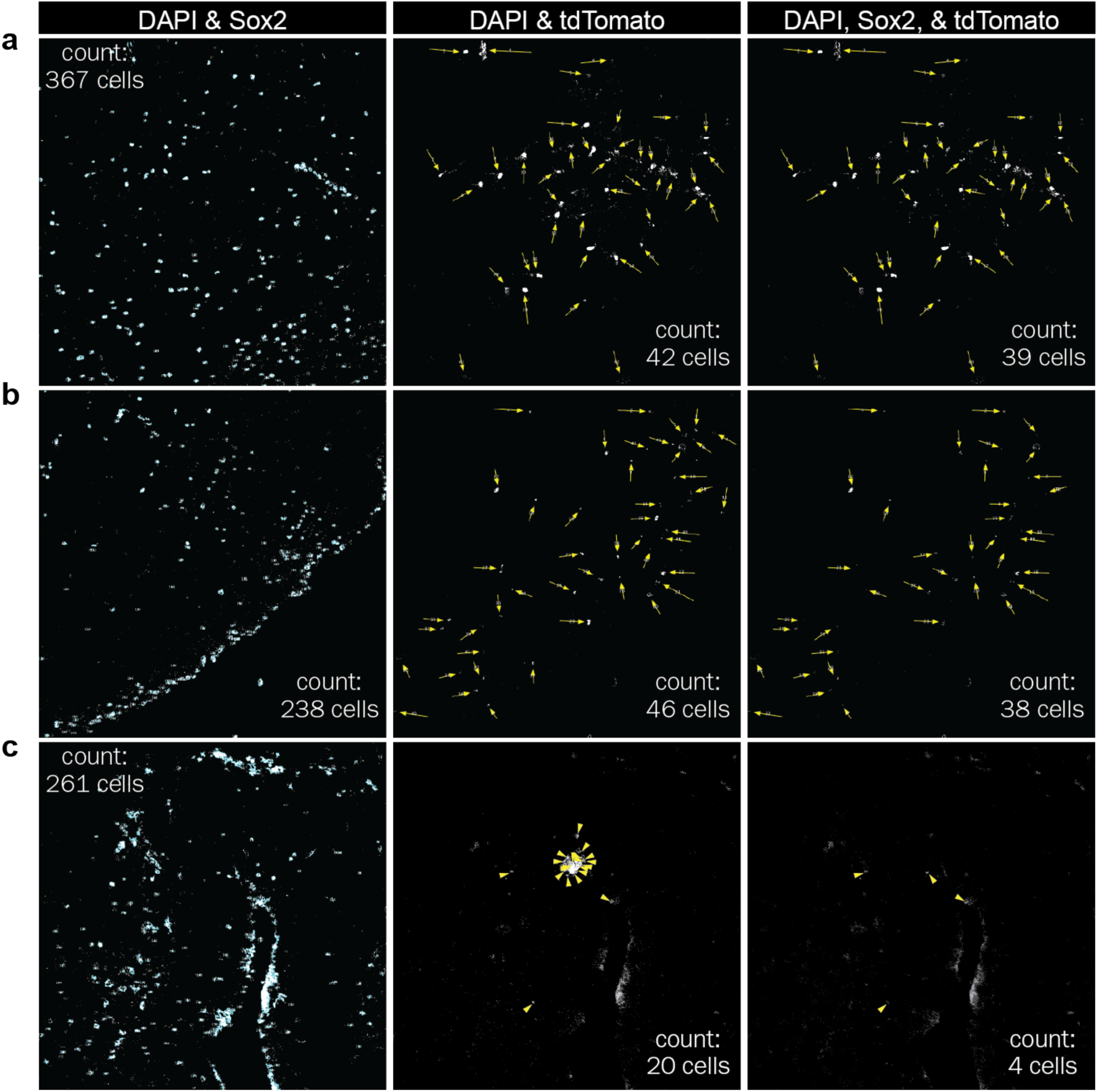
Quantification of fetal brain tissue section images. **a-c,** Quantification of displayed high-magnification images from Ai9 mouse fetuses treated with DARTs conjugated to αCD56-FITC (**a,b**) or no antibody (**c**). The images analyzed here are those shown in **Fig. 7h** (**a**), **SI Fig. 5e** (**b**), and **Fig. 7j** (**c**). FIJI was used to perform quantification, as described in **Methods**. Cells containing both DAPI and Sox2 (left), DAPI and tdTomato (middle), and all three markers (right) were counted; the numbers were subsequently compared to determine the reported target-cell specificity and the percentage of Sox2^+^ cells edited in each image. Within middle and right panels, arrows indicate each counted cell, and the total number is reported in the corner of each image.

## References

1. van Haasteren, J., Li, J., Scheideler, O. J., Murthy, N. & Schaffer, D. V. The delivery challenge: fulfilling the promise of therapeutic genome editing. Nat. Biotechnol. 38, 845–855 (2020).

2. Doudna, J. A. The promise and challenge of therapeutic genome editing. Nature 578, 229–236 (2020).

3. Raguram, A., Banskota, S. & Liu, D. R. Therapeutic in vivo delivery of gene editing agents. Cell 185, 2806–2827 (2022).

4. Wright, J. F. Manufacturing and characterizing AAV-based vectors for use in clinical studies. Gene Ther. 15, 840–848 (2008).

5. Roth, G. V., Gengaro, I. R. & Qi, L. S. Precision epigenetic editing: Technological advances, enduring challenges, and therapeutic applications. Cell Chem. Biol. 31, 1422–1446 (2024).

6. Yin, H., Kauffman, K. J. & Anderson, D. G. Delivery technologies for genome editing. Nat. Rev. Drug Discov. 16, 387–399 (2017).

7. Nelson, C. E. & Gersbach, C. A. Engineering Delivery Vehicles for Genome Editing. Annual Review of Chemical and Biomolecular Engineering vol. 7 637–662 (2016).

8. Cheng, Q. et al. Selective organ targeting (SORT) nanoparticles for tissue-specific mRNA delivery and CRISPR–Cas gene editing. Nat. Nanotechnol. 15, 313–320 (2020).

9. Kim, H. et al. Lipid nanoparticle-mediated mRNA delivery to CD34+ cells in rhesus monkeys. Nat. Biotechnol. 43, 1813–1820 (2025).

10. Hamilton, J. R. et al. In vivo human T cell engineering with enveloped delivery vehicles. Nat. Biotechnol. 10.1038/s41587-023-02085-z (2024) doi:10.1038/s41587-023-02085-z.

11. Banskota, S. et al. Engineered virus-like particles for efficient in vivo delivery of therapeutic proteins. Cell 185, 250–265.e16 (2022).

12. Nyberg, W. A. et al. In vivo site-specific engineering to reprogram T cells. Nature 652, 712–721 (2026).

13. Jung, H. et al. Virus-like particles enable targeted gene engineering and pooled CRISPR screening in primary human myeloid cells. Nat. Biotechnol. 10.1038/s41587-026-03258-2 (2026) doi:10.1038/s41587-026-03258-2.

14. Ling, S. et al. Customizable virus-like particles deliver CRISPR–Cas9 ribonucleoprotein for effective ocular neovascular and Huntington’s disease gene therapy. Nat. Nanotechnol. 20, 543–553 (2025).

15. Botchkarev, V. V. et al. In vivo gene editing of human hematopoietic stem and progenitor cells using envelope-engineered virus-like particles. Nat. Biotechnol. 10.1038/s41587-025-02915-2 (2025) doi:10.1038/s41587-025-02915-2.

16. Ngo, W., et al. Mechanism-guided engineering of a minimal biological particle for genome editing. Proc. Natl. Acad. Sci. 122, e2413519121 (2025).

17. Xu, D., et al. Programmable epigenome editing by transient delivery of CRISPR epigenome editor ribonucleoproteins. Nat. Commun. 16, 7948 (2025).

18. An, M. et al. Engineered virus-like particles for transient delivery of prime editor ribonucleoprotein complexes in vivo. Nat. Biotechnol. 10.1038/s41587-023-02078-y (2024) doi:10.1038/s41587-023-02078-y.

19. Geilenkeuser, J., et al. Engineered nucleocytosolic vehicles for loading of programmable editors. Cell 188, 2637–2655.e31 (2025).

20. Mangeot, P. E. et al. Genome editing in primary cells and in vivo using viral-derived Nanoblades loaded with Cas9-sgRNA ribonucleoproteins. Nat. Commun. 10, 45 (2019).

21. Raguram, A., An, M., Chen, P. Z. & Liu, D. R. Directed evolution of engineered virus-like particles with improved production and transduction efficiencies. Nat. Biotechnol. 43, 1635–1647 (2025).

22. Yin, D. et al. Dendritic-cell-targeting virus-like particles as potent mRNA vaccine carriers. Nat. Biomed. Eng. 9, 185–200 (2025).

23. Hindi, S. M. et al. Enveloped viruses pseudotyped with mammalian myogenic cell fusogens target skeletal muscle for gene delivery. Cell 186, 2062–2077.e17 (2023).

24. Andorko, J. I. et al. Targeted in vivo delivery of genetic medicines utilizing an engineered lentiviral vector platform results in CAR T and NK cell generation. Mol. Ther. 33, 4937–4952 (2025).

25. Dobson, C. S. et al. Antigen identification and high-throughput interaction mapping by reprogramming viral entry. Nat. Methods 19, 449–460 (2022).

26. Yu, B. et al. Engineered cell entry links receptor biology with single-cell genomics. Cell 185, 4904–4920.e22 (2022).

27. Arduini, A., Katiyar, H. & Liang, C. Progress in Pseudotyping Lentiviral Vectors Towards Cell-Specific Gene Delivery In Vivo. Viruses 17, (2025).

28. Zhou, Q. et al. T-cell receptor gene transfer exclusively to human CD8+ cells enhances tumor cell killing. Blood 120, 4334–4342 (2012).

29. Joglekar, A. V. & Sandoval, S. Pseudotyped Lentiviral Vectors: One Vector, Many Guises. Hum. Gene Ther. Methods 28, 291–301 (2017).

30. Strebinger, D. et al. Cell type-specific delivery by modular envelope design. Nat. Commun. 14, 5141 (2023).

31. Hirano, M. et al. Highly Efficient Retrograde Gene Transfer into Motor Neurons by a Lentiviral Vector Pseudotyped with Fusion Glycoprotein. PLOS ONE 8, e75896 (2013).

32. Schoderboeck, L. et al. Chimeric rabies SADB19-VSVg-pseudotyped lentiviral vectors mediate long-range retrograde transduction from the mouse spinal cord. Gene Ther. 22, 357–364 (2015).

33. Shigeki Kato et al. Selective Neural Pathway Targeting Reveals Key Roles of Thalamostriatal Projection in the Control of Visual Discrimination. J. Neurosci. 31, 17169 (2011).

34. Matuszek, Z., Brown, B. L., Yrigollen, C. M., Keiser, M. S. & Davidson, B. L. Current trends in gene therapy to treat inherited disorders of the brain. Mol. Ther. 33, 1988–2014 (2025).

35. Nikolic, J. et al. Structural basis for the recognition of LDL-receptor family members by VSV glycoprotein. Nat. Commun. 9, 1029 (2018).

36. Vaughan, T. J. et al. Human Antibodies with Sub-nanomolar Affinities Isolated from a Large Non-immunized Phage Display Library. Nat. Biotechnol. 14, 309–314 (1996).

37. Honegger, A., Spinelli, S., Cambillau, C. & Plückthun, A. A mutation designed to alter crystal packing permits structural analysis of a tight-binding fluorescein–scFv complex. Protein Sci. 14, 2537–2549 (2005).

38. Lim, K. H., Huang, H., Pralle, A. & Park, S. Stable, high-affinity streptavidin monomer for protein labeling and monovalent biotin detection. Biotechnol. Bioeng. 110, 57–67 (2013).

39. Jayaraman, J. et al. CAR-T design: Elements and their synergistic function. EBioMedicine 58, 102931 (2020).

40. Fujiwara, K. et al. Hinge and Transmembrane Domains of Chimeric Antigen Receptor Regulate Receptor Expression and Signaling Threshold. Cells 9, (2020).

41. Li, N. et al. The IgG4 hinge with CD28 transmembrane domain improves VHH-based CAR T cells targeting a membrane-distal epitope of GPC1 in pancreatic cancer. Nat. Commun. 14, 1986 (2023).

42. Yang, X. et al. Comprehensive Analysis of the Therapeutic IgG4 Antibody Pembrolizumab: Hinge Modification Blocks Half Molecule Exchange In Vitro and In Vivo. J. Pharm. Sci. 104, 4002–4014 (2015).

43. Bimbo, J. F. et al. T cell-specific non-viral DNA delivery and in vivo CAR-T generation using targeted lipid nanoparticles. J. Immunother. Cancer 13, e011759 (2025).

44. Ross, G. M. S., Filippini, D., Nielen, M. W. F. & Salentijn, G. IJ. Unraveling the Hook Effect: A Comprehensive Study of High Antigen Concentration Effects in Sandwich Lateral Flow Immunoassays. Anal. Chem. 92, 15587–15595 (2020).

45. Freed, E. O. HIV-1 Gag Proteins: Diverse Functions in the Virus Life Cycle. Virology 251, 1–15 (1998).

46. Potempa, M. et al. HIV-1 Protease Uses Bi-Specific S2/S2′ Subsites to Optimize Cleavage of Two Classes of Target Sites. J. Mol. Biol. 430, 5182–5195 (2018).

47. Lee, S.-K., Potempa, M. & Swanstrom, R. The Choreography of HIV-1 Proteolytic Processing and Virion Assembly *. J. Biol. Chem. 287, 40867–40874 (2012).

48. Baird, H. A. et al. Monitoring processed, mature Human Immunodeficiency Virus type 1 particles immediately following treatment with a protease inhibitor-containing treatment regimen. AIDS Res. Ther. 2, 2 (2005).

49. Banushi, B., Joseph, S. R., Lum, B., Lee, J. J. & Simpson, F. Endocytosis in cancer and cancer therapy. Nat. Rev. Cancer 23, 450–473 (2023).

50. Lemma, E. Y., Letian, A., Altorki, N. K. & McGraw, T. E. Regulation of PD-L1 Trafficking from Synthesis to Degradation. Cancer Immunol. Res. 11, 866–874 (2023).

51. Hendrik Schmidt, J., et al. Constitutive internalization across therapeutically targeted GPCRs correlates with constitutive activity. Basic Clin. Pharmacol. Toxicol. 126, 116–121 (2020).

52. Lampe Marko et al. Dual single-scission event analysis of constitutive transferrin receptor (TfR) endocytosis and ligand-triggered β2-adrenergic receptor (β2AR) or Mu-opioid receptor (MOR) endocytosis. Mol. Biol. Cell 25, 3070–3080 (2014).

53. Lund, K. A., Opresko, L. K., Starbuck, C., Walsh, B. J. & Wiley, H. S. Quantitative analysis of the endocytic system involved in hormone-induced receptor internalization. J. Biol. Chem. 265, 15713–15723 (1990).

54. Madisen, L. et al. A robust and high-throughput Cre reporting and characterization system for the whole mouse brain. Nat. Neurosci. 13, 133–140 (2010).

55. Staahl, B. T. et al. Efficient genome editing in the mouse brain by local delivery of engineered Cas9 ribonucleoprotein complexes. Nat. Biotechnol. 35, 431–434 (2017).

56. Schmid, B. et al. Generation of two gene edited iPSC-lines carrying a DOX-inducible NGN2 expression cassette with and without GFP in the AAVS1 locus. Stem Cell Res. 52, 102240 (2021).

57. Lin, H.-C. et al. NGN2 induces diverse neuron types from human pluripotency. Stem Cell Rep. 16, 2118–2127 (2021).

58. Guibinga, G. H., Miyanohara, A., Esko, J. D. & Friedmann, T. Cell Surface Heparan Sulfate Is a Receptor for Attachment of Envelope Protein-Free Retrovirus-like Particles and VSV-G Pseudotyped MLV-Derived Retrovirus Vectors to Target Cells. Mol. Ther. 5, 538–546 (2002).

59. Macia, E. et al. Dynasore, a Cell-Permeable Inhibitor of Dynamin. Dev. Cell 10, 839–850 (2006).

60. Cureton, D. K., Massol, R. H., Saffarian, S., Kirchhausen, T. L. & Whelan, S. P. J. Vesicular Stomatitis Virus Enters Cells through Vesicles Incompletely Coated with Clathrin That Depend upon Actin for Internalization. PLOS Pathog. 5, e1000394 (2009).

61. Koivusalo, M. et al. Amiloride inhibits macropinocytosis by lowering submembranous pH and preventing Rac1 and Cdc42 signaling. J. Cell Biol. 188, 547–563 (2010).

62. Aravamudhan, P. et al. Reovirus uses macropinocytosis-mediated entry and fast axonal transport to infect neurons. PLOS Pathog. 16, e1008380 (2020).

63. Zhang, Q. et al. Heparan sulfate assists SARS-CoV-2 in cell entry and can be targeted by approved drugs in vitro. Cell Discov. 6, 80 (2020).

64. Liu, L. et al. Heparan Sulfate Proteoglycans as Attachment Factor for SARS-CoV-2. ACS Cent. Sci. 7, 1009–1018 (2021).

65. Cagno, V., Tseligka, E. D., Jones, S. T. & Tapparel, C. Heparan Sulfate Proteoglycans and Viral Attachment: True Receptors or Adaptation Bias? Viruses 11, 596 (2019).

66. Chen, X. et al. Programmable macromolecule delivery via engineered trogocytosis. Nat. Cell Biol. 28, 1076–1086 (2026).

67. Candelaria, P. V., Leoh, L. S., Penichet, M. L. & Daniels-Wells, T. R. Antibodies Targeting the Transferrin Receptor 1 (TfR1) as Direct Anti-cancer Agents. Front. Immunol. Volume 12-2021, (2021).

68. Neil Stuart, Martin Francisco, Ikeda Yasuhiro, & Collins Mary. Postentry Restriction to Human Immunodeficiency Virus-Based Vector Transduction in Human Monocytes. J. Virol. 75, 5448–5456 (2001).

69. Ling, K.-H. et al. Deep sequencing analysis of the developing mouse brain reveals a novel microRNA. BMC Genomics 12, 176 (2011).

70. Zhang, S., Zhao, J., Zhao, C., Su, L. & Jiao, J. BCAT1 controls embryonic neural stem cells proliferation and differentiation in the upper layer neurons. Mol. Brain 16, 53 (2023).

71. Gao, K. et al. Widespread Gene Editing in the Brain via In Utero Delivery of mRNA Using Acid-Degradable Lipid Nanoparticles. ACS Nano 18, 30293–30306 (2024).

72. Brazel, C. Y. et al. Sox2 expression defines a heterogeneous population of neurosphere-forming cells in the adult murine brain. Aging Cell 4, 197–207 (2005).

73. Panciera, T. et al. Induction of Expandable Tissue-Specific Stem/Progenitor Cells through Transient Expression of YAP/TAZ. Cell Stem Cell 19, 725–737 (2016).

74. Baldauf, C. et al. Murine maternal dietary restriction affects neural Humanin expression and cellular profile. J. Neurosci. Res. 98, 902–920 (2020).

75. Walter, R. B., Boyle, K. M., Appelbaum, F. R., Bernstein, I. D. & Pagel, J. M. Simultaneously targeting CD45 significantly increases cytotoxicity of the anti-CD33 immunoconjugate, gemtuzumab ozogamicin, against acute myeloid leukemia (AML) cells and improves survival of mice bearing human AML xenografts. Blood 111, 4813–4816 (2008).

76. van der Jagt, R. H. C. et al. Localization of Radiolabeled Antimyeloid Antibodies in a Human Acute Leukemia Xenograft Tumor Model1. Cancer Res. 52, 89–94 (1992).

77. Rurik, J. G. et al. CAR T cells produced in vivo to treat cardiac injury. Science 375, 91–96 (2022).

78. Billingsley, M. M. et al. In Vivo mRNA CAR T Cell Engineering via Targeted Ionizable Lipid Nanoparticles with Extrahepatic Tropism. Small 20, 2304378 (2024).

79. Tombácz, I. et al. Highly efficient CD4+ T cell targeting and genetic recombination using engineered CD4+ cell-homing mRNA-LNPs. Mol. Ther. 29, 3293–3304 (2021).

80. Nooraei, S. et al. Virus-like particles: preparation, immunogenicity and their roles as nanovaccines and drug nanocarriers. J. Nanobiotechnology 19, 59 (2021).

81. DePolo, N. J. et al. VSV-G Pseudotyped Lentiviral Vector Particles Produced in Human Cells Are Inactivated by Human Serum. Mol. Ther. 2, 218–222 (2000).

82. Glascock, J. J. et al. Delivery of Therapeutic Agents Through Intracerebroventricular (ICV) and Intravenous (IV) Injection in Mice. J. Vis. Exp. e2968 (2011) doi:10.3791/2968.

83. Gao, K. et al. In utero delivery of mRNA to the heart, diaphragm and muscle with lipid nanoparticles. Bioact. Mater. 25, 387–398 (2023).

84. Kumar, P. et al. In Utero Transplantation of Placenta-Derived Mesenchymal Stromal Cells for Potential Fetal Treatment of Hemophilia A. Cell Transplant. 27, 130–139 (2018).

